# Ganglioside GM1-enriched rafts regulate the neuronal chloride co-transporter KCC2

**DOI:** 10.64898/2026.03.22.713396

**Authors:** Cem Karakus, Aliénor Passerat de la Chapelle, Anaïs Aulas, Elizaveta Boiko, Ophélie Aubry, Marion Russeau, Anais Fougou, Alice Trahin, Seija Legas, Jérémy Aubain, Florence Molinari, Sabine Lévi, Claudio Rivera, Coralie Di Scala

## Abstract

During brain development, dynamic remodeling of membrane lipid composition accompanies the maturation of inhibitory neurotransmission and the progressive establishment of low intracellular chloride levels. Central to this developmental transition is the neuronal K⁺–Cl⁻ cotransporter KCC2, whose stabilization at the plasma membrane enables the emergence of hyperpolarizing GABAergic signaling. Although KCC2 regulation by protein partners has been extensively characterized, whether lipid remodeling actively contributes to its membrane organization and chloride transport remains unclear. Here we identify the ganglioside GM1, a complex lipid abundant in plasma membrane of neurons, as a developmentally regulated lipid determinant of KCC2 membrane localization and function. We show that KCC2 interacts with GM1 within plasma membrane lipid rafts and that this interaction increases during postnatal brain maturation. Molecular modeling identified a conserved ganglioside-binding domain (GBD) in KCC2 centered on tryptophan 318 (W318). Biophysical analyses revealed a specific and saturable interaction between this domain and GM1 that is abolished by the epilepsy-associated W318S mutation. Disruption of KCC2–GM1 interactions, either by W318S mutation or by pharmacological depletion of GM1, excludes KCC2 from lipid rafts, alters its membrane diffusion and clustering, and reduces its surface stability. Functionally, these perturbations impair KCC2-mediated chloride extrusion and disrupt the somato-dendritic chloride gradient in hippocampal neurons. Consistent with these cellular effects, GM1-deficient (St3gal5⁻/⁻) mice exhibit selective reduced hippocampal KCC2 expression. Together, these findings reveal a lipid–protein mechanism that links developmental membrane remodeling to KCC2 stabilization and chloride homeostasis, highlighting membrane lipids as active regulators of transporter maturation and inhibitory circuit development.

## Introduction

The neuronal plasma membrane is a highly specialized molecular environment in which lipid composition plays a central role in synaptic signaling and neuronal excitability^1,2^. Far from only serving as passive structural elements, lipids, which constitute the majority of the brain’s dry mass^3,4^, regulate membrane protein trafficking^5^, conformational states and nanoscale organization^6^. Accordingly, postnatal brain development is accompanied by extensive lipid remodeling and gives rise to membrane microdomains that enable compartmentalized signaling and neuronal maturation^7,8^. Among these lipids, gangliosides are especially enriched in neurons, accounting for up to 10% of the total brain lipids^9,10^. Their abundance is tightly regulated across brain development and maturation^11^, and they are essential for normal brain function^12,13^. Genetic disruption of ganglioside biosynthesis leads to profound neurodevelopmental disorders, including epilepsy and cognitive impairment^14,15,16,17^. Gangliosides preferentially partition into cholesterol-rich lipid rafts^18,19^, nanodomains that organize receptors, ion channels, and signaling complexes^20^, suggesting a potential role in the spatial coordination of membrane signaling. Yet, despite their abundance and clinical relevance, whether gangliosides directly regulate membrane proteins that control neuronal excitability remains largely unexplored.

One of the membrane proteins, in which such ganglioside-dependent regulation could be particularly impactful, is the neuron-specific K⁺–Cl⁻ cotransporter KCC2, a key determinant of inhibitory synaptic transmission. In mature neurons, fast inhibitory synaptic transmission is mediated by GABA_A_ receptors (GABA_A_Rs) and glycine receptors (GlyRs) and critically depends on the intracellular chloride concentration [Cl^−^]i. Chloride homeostasis thus governs both the polarity and amplitude of Cl^-^ fluxes through GABA_A_R and GlyR, preserving the proper inhibitory synaptic efficacy^21^. In mature neurons, synaptic responses mediated by these receptors are hyperpolarizing, a property that emerges during brain maturation through the upregulation of the neuronal K^+^-Cl^-^ co-transporter (KCC2) which extrudes Cl^-^ from neurons under resting conditions^22^. Conversely, disruption of KCC2 functional expression compromises chloride homeostasis and is strongly associated with aberrant network activity in multiple neurological disorders, including epilepsy^23,24,25^.

Beyond developmental transcriptional and post-translational regulation, KCC2 function is dynamically controlled at the plasma membrane. Neuronal activity modulates its stability, lateral diffusion, and transport efficacy through phosphorylation-dependent mechanisms^26,27,28, 29^, together with structural interactions with other membrane proteins^30^ and the actin cytoskeleton^31,32^. These findings highlight the importance of membrane organization for KCC2 function. However, the contribution of the lipid environment itself, particularly ganglioside raft components, to KCC2 regulation remains poorly understood.

In this study, we investigated whether a lipid raft enriched ganglioside, the monosialotetrahexosylganglioside (GM1), modulates KCC2 membrane expression, dynamics, and function in mature neurons. We show that GM1 directly interacts with KCC2, stabilizing it within lipid raft domains. Disruption of this interaction, either by pharmacological depletion of membrane GM1 or by a KCC2 point mutation linked to a human neurodevelopmental disorder^33^, reduces KCC2 surface expression, alters its membrane diffusion behavior, clustering and impairs chloride extrusion. These findings uncover a previously unrecognized ganglioside-dependent mechanism that tunes neuronal chloride homeostasis and GABAergic inhibition. By identifying GM1 as a key regulator of KCC2 functional expression, our work reveals a new layer of inhibitory control and suggests that targeting lipid–protein interactions may represent a novel therapeutic strategy for disorders linked to impaired chloride homeostasis, such as epilepsy.

## Results

### KCC2 is expressed in lipid rafts

Given GM1’s potential ability to regulate GPCR^34^ and neurotransmitters receptors^35^ function, we asked whether GM1 might similarly influence KCC2 and what would be the underlying mechanism. We first investigated KCC2 and GM1 spatial and temporal distribution within neurons. Using an immunocytochemistry approach in cultured hippocampal neurons (DIV3-21), we assessed KCC2 and GM1 distribution during neuronal maturation, a period when functional KCC2 expression increases alongside with the development of GABA_A_R-mediated hyperpolarizing and inhibitory responses^36^. We showed that both KCC2 and GM1 levels progressively increased during the neuronal maturation in cell cultures from 0.134 ± 0.007 a.u. at DIV3 to 0.575 ± 0.042 a.u. at DIV 21 for KCC2 (p < 0.0001) and from 0.221 ± 0.009 a.u. to 0.517 ± 0.033 a.u. for GM1 (p < 0.0001; Fig. 1a-b). Both molecules display a punctate distribution within the somatodentritic compartment (Fig. 1a-b, Supplementary Fig. 1a). GM1 immunoreactivity was restricted to neurons Fig. 1a, Supplementary Fig. 1a) and barely detectable in astrocytes (Fig. 1a, Supplementary Fig. 1a, Supplementary Fig. 2a). Quantitative co-localization analyses showed significant developmental increases in the Pearson’s coefficient and in mutual overlap of KCC2 and GM1 over DIV in neurons (Supplementary Fig. 1a-d). Pearson’s coefficient went from 0.562 ± 0.027 at DIV3 to 0.786 ± 0.017 at DIV21 in neurons (p < 0.0001; Supplementary Fig. 1a-b). Conversely, since these two molecules were not expressed in astrocytes, no colocalization was detected in those cells (Supplementary Fig. 2a-b). This indicates that these two molecules exhibit a concurrent upregulation in neurons and spatial association during neuronal maturation.

**Figure 1:**
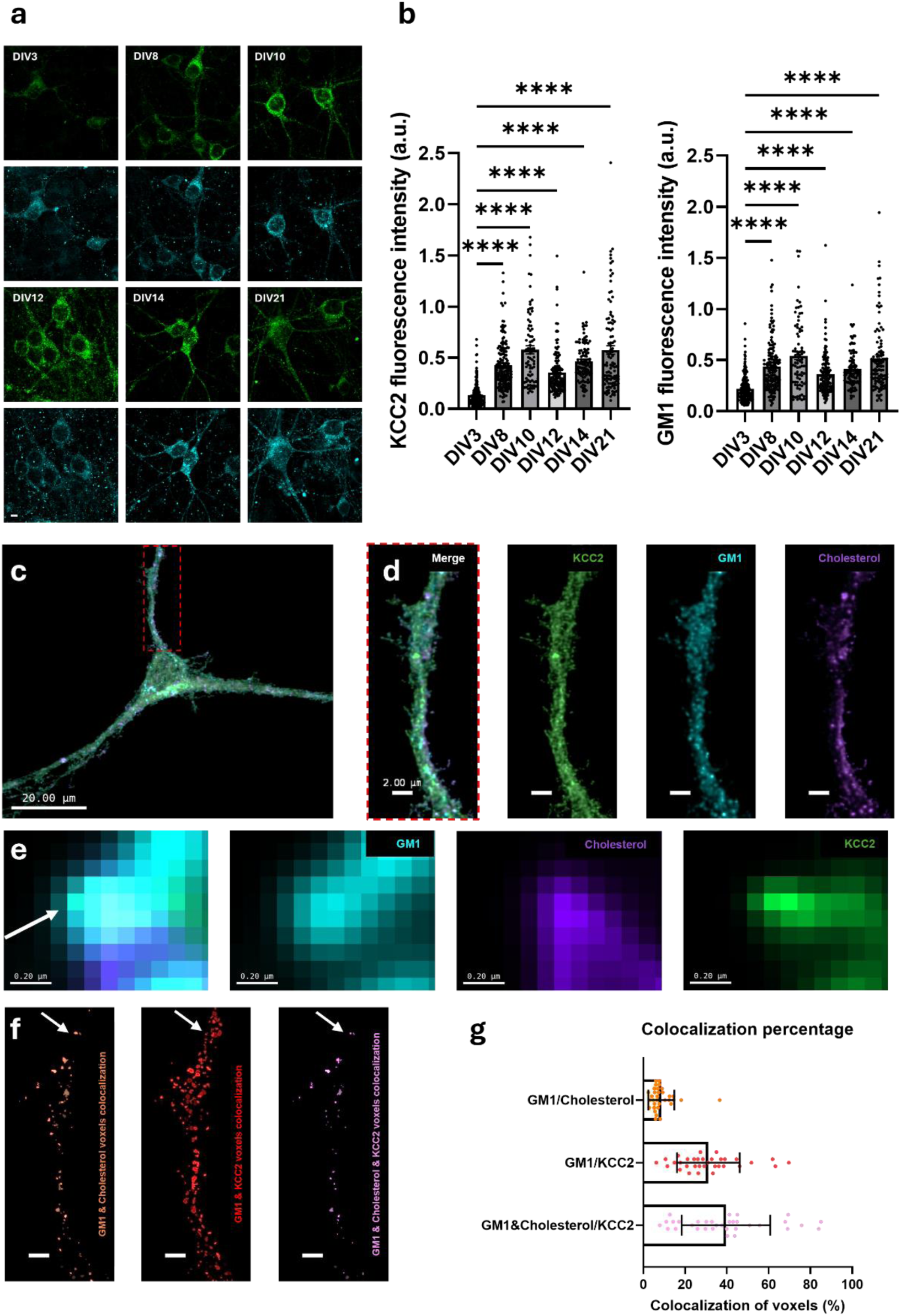
GM1, Cholesterol, and KCC2 colocalize in plasma membrane of neurons. **a.** Representative confocal images of primary hippocampal neuron cultures at DIV3-21, stained with the KCC2 (green) and GM1 (blue) antibodies. Scale bar 5 µm. **b.** Quantification of KCC2 (left) and GM1 (right) staining from pictures in **a**, data expressed as mean ± SEM, three independent experiments, Kruskal-Wallis test, ****p < 0.0001. **c.** Representative confocal images of primary hippocampal neuron cultures at DIV14, stained with the cholesterol probe (purple) and antibodies against GM1 (blue) and KCC2 (green). Scale bar: 20 µm. **d-e.** Zoomed different channel configuration of the panel **c**, showing colocalization (arrow) between cholesterol, GM1 and KCC2. Scale bar: 2 µm (**d**) and 0.2 µm (**e**). **f.** Artificial channel showing the colocalized pixel in the different configuration. Scale bar: 2 µm. **g.** Quantification of colocalization between cholesterol, GM1 and/or KCC2. All Pictures are representatives from three independent experiments made in duplicate, between 7 and 8 neurons were analyzed for each batch; for each neuron, 1–2 primary branches were quantified.

Because GM1 is enriched in cholesterol-rich lipid rafts^37^, which can modulate KCC2 functional expression^38,39^, we next examined whether KCC2 may localize in these microdomains. We used the His₆-mCherry-tagged perfringolysin theta toxin D4 domain as a raft-cholesterol probe^40^, together with GM1 and KCC2 staining at DIV14 hippocampal neurons, when KCC2 expression is maximal^36^. At this stage, cholesterol exhibited punctate distributions along the plasma membrane (Fig. 1c-f; Supplementary Fig. 3). Co-localization analysis revealed that 8.58% ± 6.19% of raft-cholesterol overlapped with GM1 (Fig. 1f-g; Supplementary Fig. 3), 31.07% ± 15.03% of GM1 puncta overlapped with KCC2 (Fig. 1f-g; Supplementary Fig. 3) and 39.53% ± 21.21% of KCC2 puncta co-localized with both GM1 and raft cholesterol (Fig. 1f-g; Supplementary Fig. 3). These data indicate that KCC2 is in GM1-containing lipid rafts, establishing a spatial framework to investigate the molecular basis of KCC2-GM1 association.

### KCC2 interacts with ganglioside GM1

Since both KCC2 and GM1 are localized into lipid rafts, we next asked whether these two molecules interact within the plasma membrane of neurons. Because lipid-protein interactions from intact plasma membranes are technically challenging to assess, we first optimized a lipid-protein co-immunoprecipitation approach in a heterologous system. HEK293 cells were transfected to overexpress WT-KCC2-mCherry (Fig. 2a), a construct previously showed to reproduce the distribution and functional properties of endogenous KCC2 in neurons^41^. In this system, immunoprecipitation with an anti-GM1 antibody pulled down KCC2 (Fig. 2b-c), indicating that the two molecules reside in the same membrane complex. Reciprocal experiments, in which KCC2 was immunoprecipitated and GM1 detected by dot blot, confirmed this association (Fig. 2b-c).

**Figure 2:**
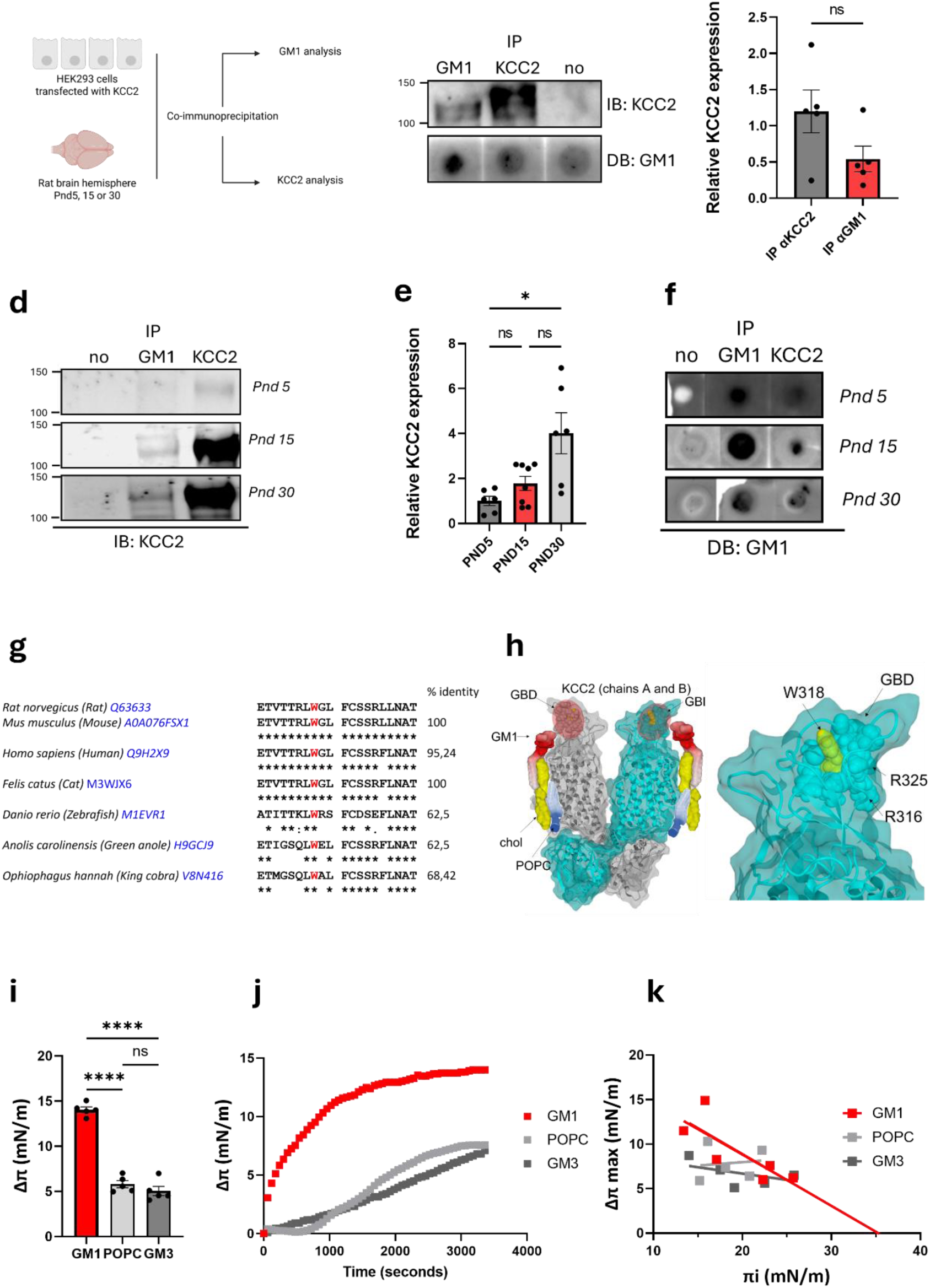
GM1 and KCC2 interact. **a.** Experimental approach of samples preparation from transfected HEK293 cells and brain rats. **b-c.** Western blot (IB), dotblot (DB) **b** and quantification of WT-KCC2-mCherry overexpressed in HEK293 cells after immunoprecipitation (IP) with a KCC2 antibody (dark grey) or a GM1 antibody (red) (**c**, mean ± SEM, five independent experiments, Welch’s test ns – not significant). **d-f.** Western blot (IB) **d**, quantification of KCC2 at PND 5 (dark grey), PND15 (red) and PND30 (light grey) after immunoprecipitation with a GM1 antibody or a KCC2 antibody **e** (mean ± SEM, n = 6-8, ANOVA test, ns – not significant *p < 0.05), dotblot (DB) of GM1 after immunoprecipitation with a GM1 antibody or a KCC2 antibody **f**. **g.** The sequence alignment depicts amino acids localized in the third extracellular loop of KCC2 from *Rattus norvegicus*; *Mus musculus*; *Homo sapiens*; *Anolis carolinensis*; *Danio rerio*. The ganglioside binding domain (GBD) is present in all sequences, centered on the tryptophane 318 (in red) that is conserved in all vertebrates and framed by basic amino acids. **h.** Molecular dynamics simulation of KCC2 dimer (grey and cyan) in presence of GM1 (red), POPC (dark blue) and cholesterol (yellow). The circle in red showed the GBD area that interacts with GM1 (left panel). A zoom on the GBD area revealed W318 as critical to structure the GBD and sustain the interaction with GM1. **i.** Structure-activity studies of WT-KCC2-GBD interactions with GM1 (red), POPC (light grey) or GM3 (dark grey) monolayer (mean ± SEM, n = 5, ANOVA test, ****p < 0.0001). **j.** Kinetic studies of WT-KCC2-GBD interaction with monolayers of GM1 (red), POPC (light grey) or GM3 (dark grey) monolayer. **k.** Specificity of WT-KCC2-GBD interactions with GM1 (red), POPC (light grey) or GM3 (dark grey) monolayer.

During brain development, KCC2 expression increases and contributes to the maturation of GABA_A_R transmission^22^. We therefore examined whether the association between KCC2 and GM1 changes over postnatal development during brain maturation. To validate our model, we first quantified KCC2, GM1, and cholesterol levels in rat brain hemispheres at postnatal days (PND) 5, 15, and 30. As expected, KCC2 expression increased markedly across brain maturation (0.281 ± 0.077 a.u.; 1.336 ± 0.408 a.u. and 1.444 ± 0.380 a.u. respectively; Supplementary Fig. 4a). GM1 levels also increased over this period, whereas cholesterol content remained relatively constant (Supplementary Fig. 4b-c). Lipid–protein co-immunoprecipitation revealed that anti-GM1 antibody co-immunoprecipitated KCC2 at all stages examined, with a progressive rise in the KCC2 fraction from 1.003 ± 0.409 a.u. to 4.009 ± 1.637 a.u; between PND5 and PND30 (Fig. 2d-e). Consistently, the reciprocal experiment confirmed the presence of GM1 in KCC2 immunoprecipitated at each maturation stage (Fig. 2f). Together, these results demonstrate that KCC2 and GM1 are part of the same membrane complex and that their association strengthens during brain maturation in parallel with increased KCC2 expression.

Within the plasma membrane, proteins interact with specific lipids through geometric and physicochemical complementarity^42,43^. These interactions help define protein’s structure, modulate their activity and influence their localization. Many proteins engage gangliosides through specialized lipid-binding motifs known as ganglioside-binding domains (GBDs)^44^. Bioinformatic analysis of the KCC2 primary sequences from multiple species revealed the presence of such a domain (Fig. 2g), suggesting potential interaction with gangliosides, including GM1. In both human and rodent KCC2, this GBD spans amino acid residues 316-325 within the third extracellular loop and is highly conserved across species (Fig. 2g). The domain centers on a tryptophan residue (W318), an aromatic amino acid frequently critical for interaction with gangliosides^44,45^. To test the ability of this domain to interact with GM1, we realized molecular dynamic simulation of KCC2 with GM1 molecules. We showed that the KCC2 ganglioside binding domain (KCC2-GBD) fits with the GM1 molecules (Fig. 2h).

To test the functional capacity of this domain to interact with GM1, we performed lipid monolayer experiments using the Langmuir films system. In this assay, lipid–protein interaction is indicated by insertion of the protein into the monolayer, resulting in an increase in surface pressure^46,47^ (Supplementary Fig. 5a). The KCC2-GBD produced a strong increase in the surface pressure upon injection beneath GM1 monolayers (Δπ = 14.045 ± 0.316 mN.m^-1^), demonstrating a robust interaction (Fig. 2i). The domain also interacted, though less strongly, with other neuronal gangliosides (disialoganglioside 1a (GD1a), disialoganglioside 1b (GD1b) and trisialoganglioside 1b (GT1b)) (Supplementary Fig. 6a) and only weakly with glial gangliosides like monosialodihexosylganglioside (GM3, Δπ = 5.038 ± 0.514 mN.m^-1^) or with the main lipid of the plasma membrane, the palmitoyloleoylphosphatidylcholine (POPC, Δπ = 5.803 ± 0.338 mN.m^-1^) that was used here as a negative control (Fig. 2i). Kinetic analyses further showed that KCC2-GBD associated with GM1 more rapidly than with other lipids (Fig. 2j; Supplementary Fig. 6b).

We next quantified the specificity of this interaction by determining the critical pressure of insertion (πc) (Fig. 2k; Supplementary Fig. 6c). Lipid monolayers were prepared at increasing initial surface pressures (πi), and the maximal surface pressure variation (Δπ) after 1 h was measured. A characteristic decrease in Δπ with increasing πi—a hallmark of specific lipid–protein interaction—was observed for GM1, but not for GM3 or POPC (Fig. 2k). From this relationship, we calculated a πc for the KCC2-GBD with GM1, representing the surface pressure above which insertion no longer occurs due to lipid packing constraints^47^. Critical pressures were also determined for GD1a, GD1b, and GT1b, confirming interaction with neuronal gangliosides in general, albeit with lower affinity than for GM1 (Supplementary Fig. 6c). No πc was detected for GM3 or POPC, indicating non-specific interactions with these lipids. Finally, dose–response experiments performed at a fixed initial surface pressure (15 ± 1 mN.m^-1^) revealed a saturable binding profile, further supporting the specificity of the KCC2-GBD–GM1 interaction (Supplementary Fig. 5b). Together, these results establish that KCC2 contains a conserved ganglioside-binding domain that specifically recognizes and interacts with GM1, defining GM1 as a key component of KCC2’s immediate lipid environment in the plasma membrane.

### A specific KCC2–GM1 interaction stabilizes KCC2 within lipid rafts

Our modeling simulation indicates that the tryptophan residue at position 318 (W318) plays a pivotal role in mediating the interaction by structuring the GBD and enabling the interaction with GM1. Interestingly, a point mutation substituting this residue with a serine (W318S) has been linked to a neurodevelopmental disease, although the functional consequences for KCC2 remain poorly understood^33^. Based on these observations, we hypothesized that W318 is critical for the KCC2-GM1 interaction. To test this hypothesis, we conducted lipid monolayer experiments using the KCC2-GBD carrying the W318S mutation (W318S-KCC2-GBD). Consistent with our model, W318S-KCC2 produced only a modest increase in surface pressure (5.204 ± 0;162 mN.m^-1^ after 1 h), compared with a 14.446 ± 0.299 mN.m^-1^ increase observed for the WT-KCC2-GBD (Fig. 3a-b). As expected, increasing the initial surface pressure of the GM1 monolayer did not attenuate the surface pressure variation, and no critical pressure of insertion could be determined for the W318S-KCC2-GBD (Fig. 3c). These findings indicate that, unlike the wild-type domain, the W318S mutant fails to insert into the GM1 monolayer.

**Figure 3:**
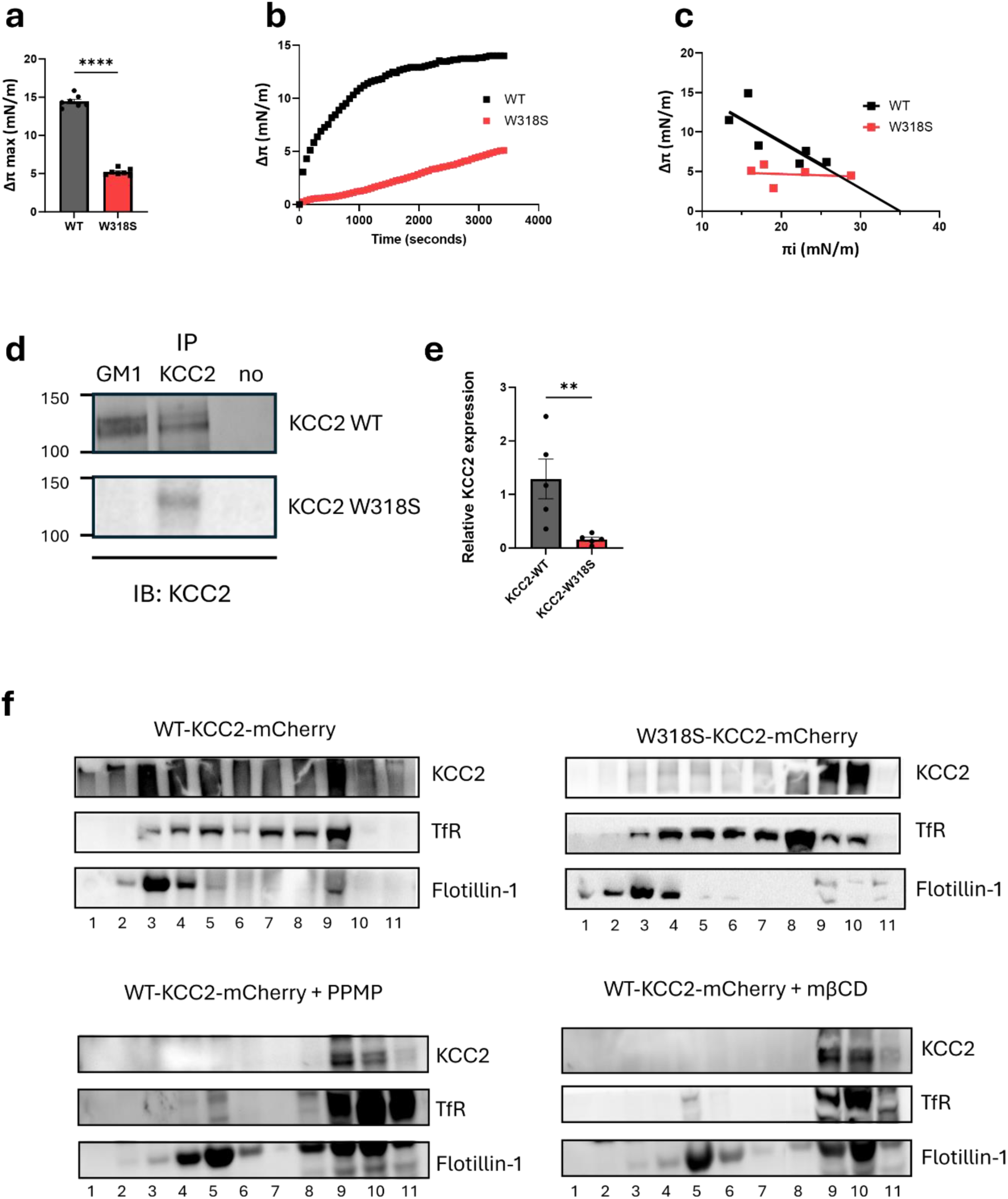
The interaction of GM1 with KCC2 is specific. **a.** Structure-activity studies of WT-KCC2-GBD (dark grey) and W318S-KCC2-GBD (red) interactions with GM1 monolayers (mean ± SEM, n = 7, t-test, ****p < 0.0001). **b.** Kinetic studies of WT-KCC2-GBD (black) and W318S-KCC2-GBD (red) interaction with GM1monolayer. **c.** Specificity of WT-KCC2-GBD (black) and W318S-KCC2-GBD (red) interactions with GM1 monolayer. **d-e.** Western blot **d** and quantification of WT-KCC2-mCherry (dark grey) or W318S-KCC2-mCherry (red) overexpressed in HEK293 cells after immunoprecipitation with a GM1 antibody (mean ± SEM, n = 5, Mann-Whitney, **p < 0.01) **e**.

To further evaluate the importance of W318 in the context of the full-length KCC2, we generated a mutant construct expressing W318S-KCC2-mCherry. We first checked that both WT-KCC2-mCherry and W318S-KCC2-mCherry displayed comparable transfection efficiency (Supplementary Fig. 7a-b). Co-immunoprecipitation assays on HEK293 cells transfected with the full-length WT-KCC2-mCherry or the full length W318S-KCC2-mCherry revealed that the anti-GM1 antibody pulled down markedly less W318S-KCC2-mCherry 0.162 ± 0.041 a.u. than WT-KCC2-mCherry 1.289 ± 0.372 a.u. (p = 0.008; Fig. 3d-e). This indicates that W318 is critical for the interaction of KCC2 with GM1.

We next examined whether the W318S mutation or lipid rafts disruption affected KCC2 localization within GM1-enriched microdomains. In sucrose gradient fractioning, WT-KCC2-mCherry, beforehand expressed in HEK293 cells, was distributed between low-density, detergent-resistant, fractions containing the raft markers flotillin-1 (fractions 3 to 6) and the detergent-soluble fractions containing the transferrin-receptor (fractions 8 to 11) (Fig. 3f). Conversely, the W318S-KCC2-mCherry mutant was largely excluded from the low-density fractions and distributed to detergent-soluble regions (fractions 9 to 11; Fig. 3f). Similarly, depletion of membrane cholesterol with methyl-β-cyclodextrin (MβCD) or depletion of membrane GM1 with 1R,2R-(+)-1-phenyl-2 palmitoylamino-3-N-morpholine-1-propanol (PPMP) that inhibits gangliosides synthesis (Supplementary Fig. 8a-c) caused a comparable shift of WT-KCC2 from raft to non-raft fractions (Fig. 3f: similar results were observed on 4 independent experiments). Together, these results demonstrate that the interaction between KCC2 and GM1, mediated by W318, is essential for the stable incorporation of KCC2 into lipid rafts and thus for its proper membrane partitioning.

### Association with GM1-containing lipid rafts promotes KCC2 stability at the plasma membrane

To assess whether the interaction with GM1 affects KCC2 stability at the plasma membrane, we performed live-staining analysis of surface and internalized KCC2 fractions using a pH-sensitive fluorescent construct (KCC2-pHext)^41^. This reporter, tagged in the second extracellular loop, enables quantitative distinction of the total (F_t_), surface-expressed (F_m_), and internalized (F_i_) KCC2 pools^48^. We compared five conditions: WT-KCC2-pHext, A/A-KCC2-pHext (positive control; increased surface expression), ΔNTD-KCC2-pHext (negative control; impaired surface trafficking), WT-KCC2-pHext after PPMP treatment, and W318S-KCC2-pHext. In these conditions, PPMP treatment was non-toxic for Neuro2a cells (Supplementary Fig. 9a) and effectively reduced GM1 membrane levels (Supplementary Fig. 9b-c). All constructs showed comparable total expression levels (F_t_: 0.674 ± 0.094; 0.555 ± 0.116; 1.122 ± 0.202; 0.549 ± 0.088 and 0.761 ± 0.111 a.u., respectively; Fig. 4a-b). As expected, surface expression was highest for the A/A mutant than WT-KCC2-pHext (F_m_ = 0.595 ± 0.088 vs 0.289 ± 0.030 a.u.) and nearly abolished for ΔNTD-KCC2 (F_m_ = 0.005 ± 0.003 a.u.), validating the assay (Fig. 4a, 4c). Reducing GM1 levels by PPMP treatment did not significantly alter the surface fraction of WT-KCC2-pHext (F_m_ = 0.175 ± 0.021 a.u. vs. 0.289 ± 0.030 a.u. for control; Fig. 4a, 4c), though it modestly increased internalization (F_i_ = 0.296 ± 0.062 a.u. for PPMP vs. 0.260 ± 0.030 for control^15^ a.u.; Fig. 4a, 4d). By contrast, the W318S mutation caused a strong reduction in both surface (F_m_ = 0.038 ± 0.006 a.u.) and internalized (F_i_ = 0.055 ± 0.006 a.u.) fractions relative to WT-KCC2 (F_m_ = 0.289 ± 0.030 a.u., F_i_ = 0.260 ± 0.030 a.u.; Fig. 4a–d). The W318S-KCC2-pHext profile closely resembled that of the ΔNTD mutant, consistent with a defect in surface stability.

**Figure 4:**
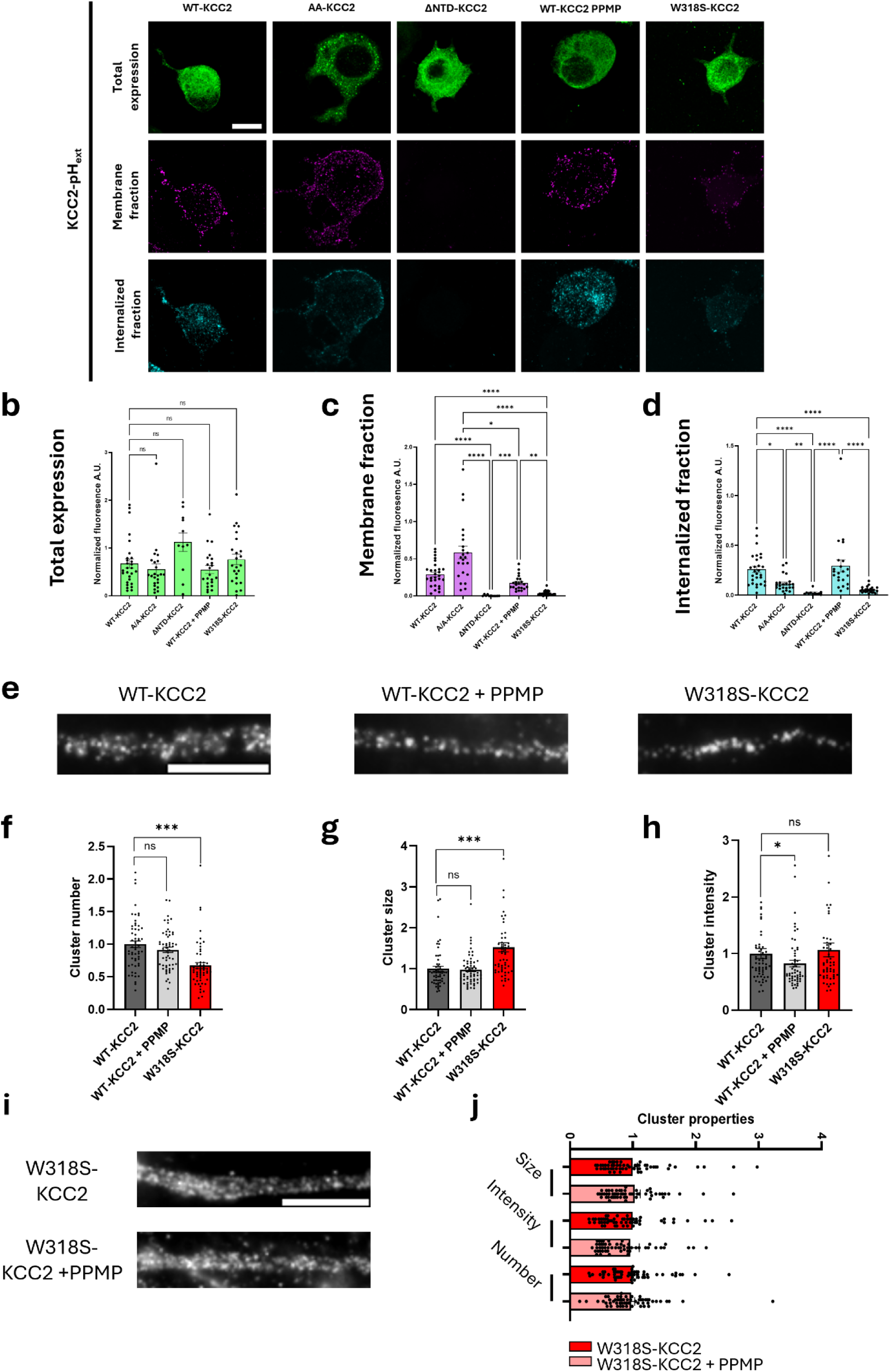
Regulation of KCC2 properties within the plasma membrane by GM1. **a.** Representative Z-stack projection of live immunostaining images of KCC2-pHext showing total expression (green), membrane fraction (cyan) and internalized fraction (magenta) for WT-KCC2 control, AA-KCC2, ΔNTD-KCC2, WT-KCC2+PPMP and W318S-KCC2. Scale bar: 10 µm. **b-d.** Quantification of KCC2-pHext total expression **b**, KCC2-pHext membrane fraction **c**, and KCC2-pHext internalized fraction **d** (mean ± SEM, three independent experiments, Kruskal–Wallis test *p < 0.05, **p < 0.01, ***p < 0.001, ****p < 0.0001, ns – not significant). **e.** Surface staining of KCC2-Flag in 21-DIV-old hippocampal neurons transfected with WT-KCC2 treated or not with 10 µM PPMP for 48 h or transfected with W318S-KCC2. Scale bar, 5 µm. **f-h.** Quantification of the mean number (**f**), size (**g**), and intensity (**h**) of KCC2 clusters for WT-KCC2 in control condition (dark grey bars) or upon application of PPMP (light grey bars) or for W318S-KCC2 (red bars). Data shown as mean ± SEM. In all graphs, values were normalized to the corresponding WT-KCC2 mean values. N=3 cultures, 55 to 58 cells. Mann-Whitney test: ns, not significant, and *** p<0.001. **i.** Surface staining of W318S-KCC2 in 21 DIV-old hippocampal neurons transfected with W318S-KCC2-Flag and treated or not with 10 µM PPMP for 48 h. Scale bar, 5 µm. **j**. Quantification of the mean number, size, and intensity of KCC2 clusters for W318S-KCC2 in control condition (red bars) or upon application of PPMP (pink bars). Data shown as mean ± SEM. In all graphs, values were normalized to the corresponding W318S-KCC2 mean values. N=3 cultures, 55 to 56 cells. Mann-Whitney test: size, p = 0.904; intensity, p = 0.287; number, p = 0.820.

We next assessed whether a change in the interaction with GM1 affects KCC2 clustering. Hippocampal neurons were transfected with Flag-tagged KCC2 (KCC2-Flag) at DIV14 and surface-labeled at DIV21 using anti-Flag antibodies. In these conditions, W318S-KCC2-Flag–expressing neurons exhibited fewer and less intense surface clusters compared to WT-KCC2-Flag (Fig. 4e-f, 4h), whereas the remaining clusters were larger in area (Fig. 4e, 4g), suggesting compensatory cluster merging. Similarly, PPMP-induced GM1 depletion on neurons (Supplementary Fig. 10a-b), reduced the intensity of WT-KCC2-Flag clusters (Fig. 4e, 4h) without affecting cluster size or number (Fig. 4f-g). Thus, PPMP treatment of WT-KCC2-Flag or expression of the W318S mutation in untreated cells reduces KCC2-Flag clustering. Importantly, combining PPMP treatment with the W318S mutation produced no additive effect (Fig. 4i-j), confirming the specificity of GM1’s influence and indicating that the mutation does not disrupt overall raft organization.

Lateral diffusion rapidelly modulates KCC2 clustering: decreased clustering correlates with higher diffusion coefficients and surface area explored, reflecting reduced membrane confinement^27,49^. We therefore investigated whether GM1 influences KCC2 lateral diffusion using quantum dot based single-particle tracking (QD-SPT). Hippocampal neurons were transfected with KCC2-Flag at DIV14 and surface-labeled at DIV21 using anti-Flag antibodies, biotinylated Fab fragments, and streptavidin-coated QDs. PPMP treatment markedly increased the membrane surface area explored by a representative KCC2 trajectory compared to control conditions (Fig. 5a).

**Figure 5:**
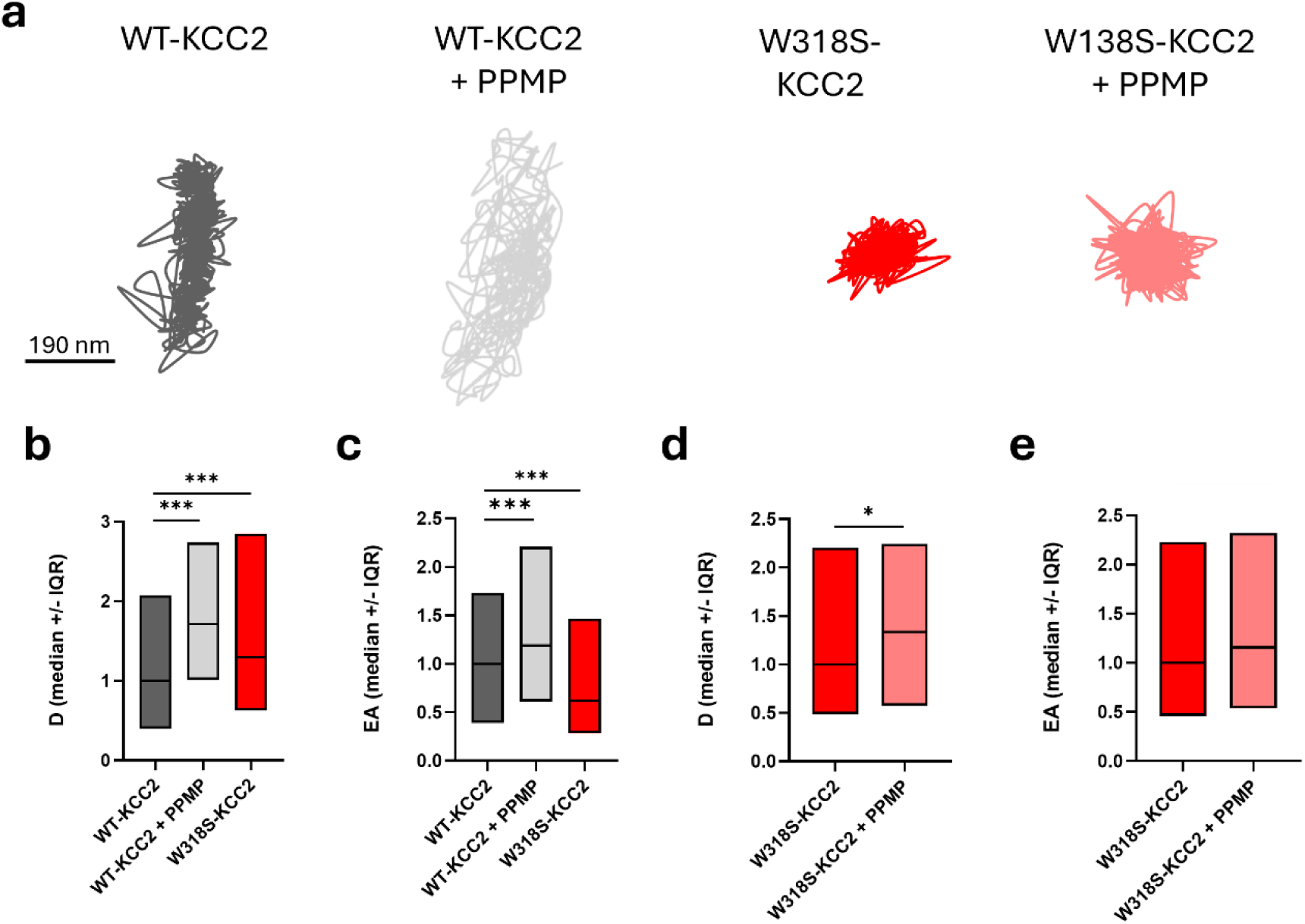
Reduced ganglioside levels or disruption of KCC2-GM1 interaction alters KCC2 lateral diffusion. **a.** Examples of trajectories for WT-KCC2 control (dark grey), WT-KCC2 + PPMP (light grey), W318S-KCC2 control (red), and W318S-KCC2 + PPMP (pink) on the surface of dendrites of mature 21-DIV-old neurons. Scale bar, 190nm. **b-c.** Median diffusion coefficient D (b) and explored area (EA) (c) ± 25–75% interquartile range (IQR) for WT-KCC2 in control condition (dark grey bars) or upon application of PPMP (light grey bars) or for W318S-KCC2 (red bars). Values were normalized to the median KCC2-WT control values. N=2 cultures, 198 to 1117 QDs. KS test *** p < 0.001. **d-e.** Median diffusion coefficient D (d) and explored area (EA) (e) ± 25–75% interquartile range (IQR) for W318S-KCC2 in control condition (red bars) or upon application of PPMP (pink bars). Values were normalized to the median W318S-KCC2 control values. KS test: d, p = 0.03; e, p = 0.07.

Analysis of a large population of QDs revealed a significant increase in both the diffusion coefficient (Fig. 5b) and the surface area explored (Fig. 5c) by WT-KCC2-Flag under PPMP treatment compared to controls, indicating increased mobility and reduced confinement, consistent with a declustering effect.

The W318S mutation of KCC2, which lacks GM1 binding, partially mimics the effect of PPMP on WT-KCC2 declustering: the diffusion coefficient of the W318S-KCC2-Flag mutant was increased compared to WT-KCC2-Flag (Fig. 5a, 5b), despite exhibiting increased confinement (Fig. 5c). This behavior generally reflects a signature of molecules targeted to endocytic pits for storage and/or internalization (ref). This hypothesis is further supported by the observation that the number of trajectories detected at the surface of neurons expressing W318S-KCC2-Flag was markedly lower than in neurons expressing WT-KCC2-Flag under the same culture and labeling conditions, indicating decreased surface expression of KCC2 (data not shown). Interestingly, PPMP treatment did not affect the confinement of W318S-KCC2-Flag and only slightly increased its diffusion coefficient (Fig. 5a, 5d–e), suggesting a weak additive effect of the treatment.

Together, these results indicate that the interaction with GM1 is essential for proper KCC2 membrane organization and stability, influencing both its diffusion dynamics and surface clustering at the neuronal plasma membrane.

### Functional impact of KCC2 regulation by GM1 lipid rafts

Disruption of KCC2 membrane organization is known to impair neuronal Cl⁻ extrusion and thereby alter GABA_A_R-mediated inhibition. We next examined the functional consequences of disrupting KCC2–GM1 interaction —either pharmacologically, using PPMP to reduce ganglioside synthesis, or genetically, with the W318S-KCC2 mutation—on KCC2 Cl⁻ transport activity. Whole-cell patch-clamp recordings were performed in primary hippocampal neurons transfected with full-length WT-KCC2 or W318S-KCC2 under control or low-GM1 conditions. Under a constant and defined intracellular Cl⁻ load via the somatic pipette, active KCC2 generates a negative somato-dendritic electrochemical Cl⁻ gradient (Fig. 6a-c)^50^. For each condition, we measured the Cl⁻ reversal potential (ECl^-^) at the soma and at proximal dendrites (∼100 µm) following brief GABA application (Fig. 6d-e) and calculated the resulting somato-dendritic gradient (Fig. 6f). Both PPMP treatment and the W318S-KCC2 mutation markedly reduced the gradient (1.467 ± 0.672 and -0.857 ± 0,273 respectively; Fig. 6f), to levels comparable to those observed in neurons expressing shRNA-KCC2 (-0.210 ± 0.482), which suppresses KCC2-mediated Cl⁻ extrusion (Fig. 6c, 6f). These results demonstrate that the interaction between KCC2 and GM1, and its proper localization within lipid rafts, are essential for maintaining efficient Cl⁻ transport and GABAergic inhibitory strength.

**Figure 6:**
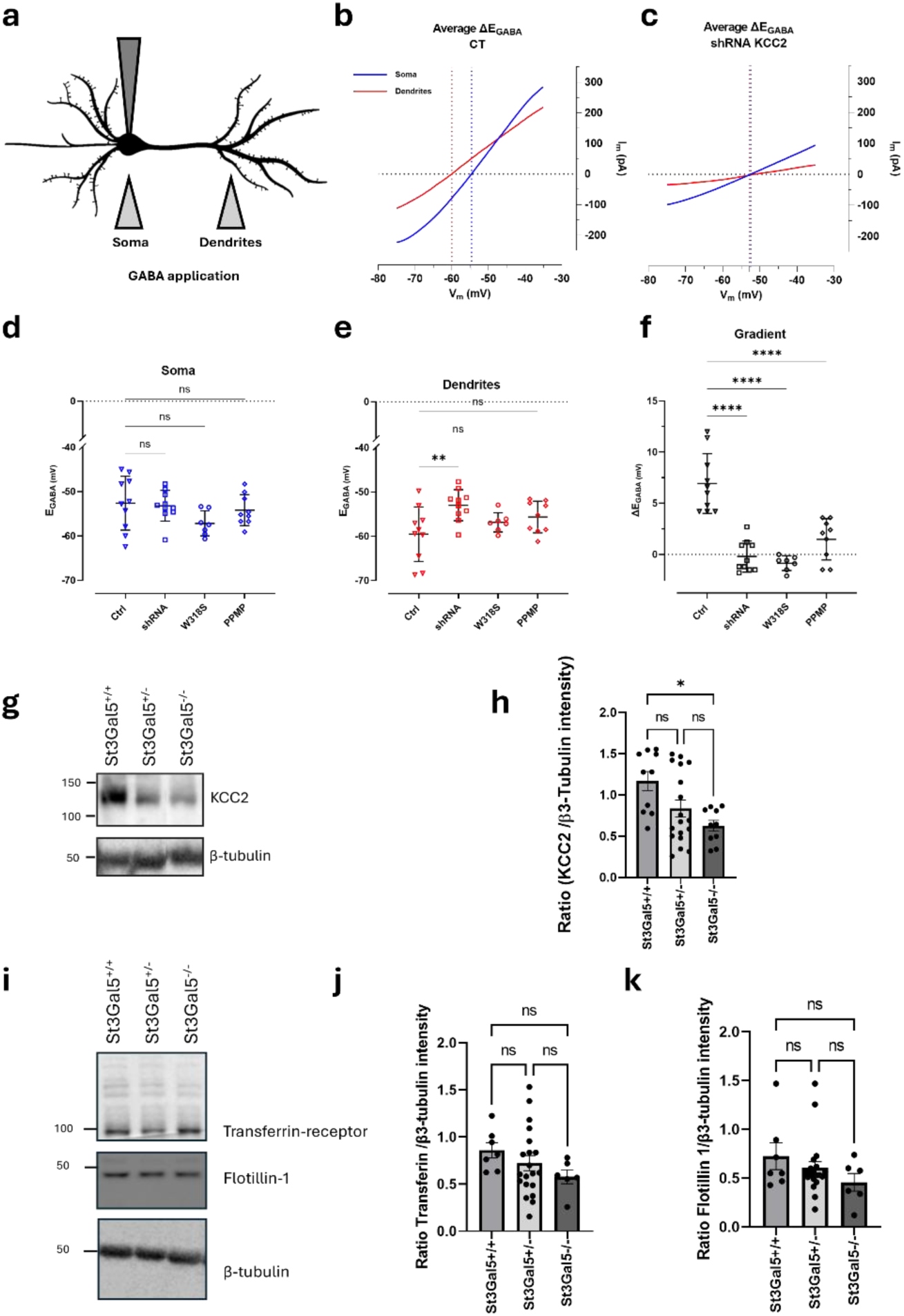
GM1 regulates KCC2 function. **a** Schematic drawing illustrating the experimental setup for assessing changes in dendritic chloride extrusion. GABA was locally applied at the soma and at the primary dendrite of a transfected neuron recorded in whole-cell voltage clamp mode with a pipette containing 19 mM of Cl^−^. **b-c** Mean voltage ramp trace during local application of GABA at the soma (blue solid line) and dendrite (red solid line), in control (CT) neurons transfected with pcDNA 3.1 (-) and eGFP **b**, and in negative control neurons transfected with shRNA against KCC2 and eGFP **c**. Red or blue dashed lines mark an estimation of the GABA_A_ reversal potential at the specific location. **d-f** E_GABA_ values at the soma **d**, dendrites **e** and ΔE_GABA_ **f** of neurons. Dots represent individual neurons, the horizontal line indicates the mean, and error bars represent SD. Normality was assessed using the Shapiro-Wilk and Kolmogorov-Smirnov tests. Group comparisons were performed using ordinary one-way ANOVA was realized to compare the groups, with **p < 0.01, ****p < 0.0001 and ns indicating not significant. **g-h.** Western blot of KCC2 and β-tubulin (**g**) and corresponding quantification (**h**) showing that GM1 loss in mouse brains led to a decrease in KCC2 expression in *St3Gal5*^+/-^ and *St3Gal5*^-/-^ compared to *St3Gal5*^+/+^ (mean ± SEM, Kruskal-Wallis, *p < 0.05). **i-k.** Western blot of transferrin-receptor, flotillin-1 and β-tubulin (**i**) and quantification (**j-k**) The loss of GM1 does not affect transferrin-receptor (**j** mean ± SEM.; ANOVA, p = n.s.) nor flotillin-1 (**k** mean ± SEM; Kruskal-Wallis, p = n.s.).

### GM1 dependent regulation of KCC2 *in vivo*

Given the strong impact of GM1 on KCC2 functional expression in cultured hippocampal neurons, we next assessed whether this mechanism operates *in vivo*. We used a genetic mouse model lacking GM1 due to ST3GAL5 deficiency^51^ (herein referred as *St3Gal5^-/-^* mice). ST3GAL5, also known as GM3 synthase or lactosylceramide α-2,3-sialyltransferase, catalyzes the first step of GM1 biosynthesis^52^. Therefore, KCC2 expressions within the hippocampus of two-month-old *St3Gal5*^+/+^, *St3Gal5*^+/-^ and *St3Gal5*^-/-^ mice were analyzed by western blot. We observed a progressive and significant reduction of KCC2 expression from 1.168 ± 0.124 a.u. on *St3Gal5*^+/+^ to 0.835 ± 0.104 a.u. and 0.627 ± 0.064 a.u. in *St3Gal5*^+/-^ and *St3Gal5*^-/-^ mice respectively (p = 0.017Fig. 6g-h). This effect was specific for KCC2, as the expression of flottilin-1 and transferrin-receptor, respectively markers of raft and non-raft membrane compartments, remained unchanged in *St3Gal5*^+/-^ and *St3Gal5*^-/-^ mice compared to *St3Gal5*^+/+^ ones (Fig. 6i-k). These results demonstrate that reduced neuronal GM1 levels lead to a selective downregulation of KCC2 expression in the hippocampus.

## Discussion

The modulation of KCC2 functional expression at the neuronal plasma membrane is essential for maintaining efficient fast inhibitory transmission^27,53^. Despite this critical role, only two studies have examined how membrane lipids—the immediate molecular environment of KCC2—influence its membrane behavior^38,39^. Although these reports reached divergent conclusions, they suggested that KCC2 partitioning between raft and non-raft plasma membrane domains modulates its functional expression. Importantly, the underlying mechanisms and their impact on KCC2 membrane behavior have remained poorly defined. Our study demonstrates that lipid rafts directly regulate KCC2 organization at the plasma membrane. We identify the GM1 ganglioside as a crucial determinant of KCC2 stabilization within raft domains, thereby modulating KCC2 localization and function. This regulation depends on a specific interaction between KCC2 and GM1 mediated by a conserved ganglioside-binding domain^44,45^ (Fig. 2g-k, Fig. 3). The KCC2–GM1 interaction emerges during neuronal maturation (Fig. 2d-f), promotes KCC2 retention at the plasma membrane (Fig. 4a-d), regulates its lateral mobility and clustering (Fig. 4e-h and Fig. 5), and supports its transporter activity (Fig. 6b-f). Conversely, disrupting this interaction alters its mobility (Fig. 5), clustering (Fig. 4e-h) and diminishes its functional activity (Fig. 6d-f). Collectively, these findings reveal a previously unrecognized lipid-dependent mechanism that governs KCC2 membrane organization and may represent a key modulatory pathway of GABAergic signaling.

KCC2 partitioning between raft and non-raft domains^38,39^ indicates that distinct subpopulations of the co-transporter coexist at the cell surface. The structural integrity of lipid rafts is essential for maintaining this organization. Consistent with this view, manipulations that reduce membrane cholesterol or ganglioside content and thereby disrupt raft architecture^54^ cause KCC2 to dissociate from raft fractions, pointing to a specific interaction between KCC2 and raft-associated lipids. Our data identify the ganglioside GM1 as the key determinant of KCC2 localization within raft domains.

To test the functional importance of this interaction, we disrupted KCC2-GM1 binding using two complementary strategies: (i) depletion of membrane GM1 by PPMP treatment (Supplementary Fig. 8-10) and (ii) expression of the GM1-binding deficient KCC2-W318S mutant (Fig. 3a-e). In both conditions, loss of KCC2–GM1 interaction led to the exclusion of KCC2 from raft domains (Fig. 3f). Consistent with Watanabe et al.^39^, the fraction of KCC2 residing outside rafts displayed reduced clustering (Fig. 4e-f) and lower transport activity (Fig. 6d-f). Together, these results show that interaction with GM1 is required to maintain KCC2 within raft domains and thereby supports both its membrane clustering (Fig. 3f) and functional activity (Fig. 6).

Mechanistically, we propose that GM1 binding stabilizes a local 3D conformation of KCC2 that favors its retention in rafts and promotes interactions required for efficient membrane organization and function. Conversely, disruption of this interaction may induce local structural rearrangements in KCC2, with secondary effects on its association with scaffolding partners and/or its ability to oligomerize. Because lipid rafts concentrate multiple signaling proteins, including kinases and phosphatases^55,56^, displacement of KCC2 from these domains is also predicted to alter upstream regulatory pathways, including WNK and SPAK/OSR1^26,57^. Relocation of KCC2 to a non-raft lipid environment may disrupt the membrane-proximal conditions required for optimal coupling to these regulatory pathways. In particular, because SPAK and OSR1 are recruited to cation-chloride cotransporters through RFXV-motif-dependent interactions^58^, and WNK kinases depend on local ionic conditions for activation, persistent localization of KCC2 outside the raft domains is likely to impair or misdirect WNK–SPAK/OSR1 signaling.

Impaired KCC2 function is a well-established contributor to neuronal hyperexcitability and epileptogenesis^59,60,61^. Our work pinpoint that changes in lipid raft KCC2 partitioning offer a plausible mechanistic link between membrane composition and seizure susceptibility. Consistent with this, ganglioside-deficient animal models—such as GM3 synthase (GM3S) knockout mice, which display altered brain GM1 levels and downstream ganglioside composition^51^—exhibit heightened neuronal excitability, aberrant hippocampal neurogenesis, and increased susceptibility to kainate- or pilocarpine-induced seizures^62,63^. In humans, GM3S deficiency leads to severe neurological dysfunction, including intractable epilepsy, intellectual disability, and blindness^12,14,15,16^, while dietary restoration of complex gangliosides ameliorates cognitive and neurogenic deficits in murine models^64^. Interestingly, functionally disabling human KCC2 mutations, except for blindness, lead to similar phenotypes^33,41,65,66^. Moreover, hippocampal tissue from patients with temporal lobe epilepsy demonstrates a selective loss of GM1 and of GD1a, a GM1-derived ganglioside, suggesting that ganglioside depletion may compromise lipid raft integrity and impair KCC2 surface stability^67^. Within this context, the W318S mutation in human KCC2, which disrupts transporter trafficking and membrane anchoring^33^, likely amplifies deficits in raft association and chloride extrusion, further promoting hyperexcitability.

Together, these results support a model in which GM1-dependent raft integrity and KCC2 structural stability act as convergent determinants of GABAergic inhibition. More broadly, GM1 ganglioside emerges as a central organizer of inhibitory membrane proteins activity. Its enrichment in lipid rafts coordinates protein-protein interactions. Consistent with this mechanism, GABA_A_R occupy GM1 / cholesterol raft domains at resting condition in hippocampal cultures but redistribute upon GABA binding, tuning receptor gating and inhibitory strength^68^. Likewise, Handlin et al. (2024) demonstrated that HCN channels in dorsal root ganglion neurons are controlled by rafts, whose disruption alters channel gating and neuronal excitability^69^. Collectively, these studies position GM1 as a key lipid determinant of neuronal excitability. By shaping raft nanodomains that spatially coordinate KCC2, GABA_A_R receptors, and other ion channels, GM1 integrates membrane architecture with synaptic signaling to maintain the excitatory–inhibitory balance critical for network stability.

## Material and methods

### Molecular Modelling

*In silico* study of KCC2 and GM1 interaction was performed with Hyperchem 8 software (ChemCAD) as previously described^70^. Briefly, the amino acid sequence of human KCC2 was retrieved from UniProt (Q9H2X9-2) and GM1 structure was obtained from CHARMM-GUI Glycolipid Modeler (http://www.charmmgui.org/?doc=input/glycolipid). Geometry optimization of KCC2 and GM1 molecules complex was achieved using Polak-Ribière algorithm. Molecular dynamics simulations were performed for iterative periods of time of 1 nsec *in vacuo* with the Bio+ (CHARMM) force field. The molecules were visualized with Hyperchem 8, PDB-viewer and Molegro Molecular Viewer softwares. The energies of interaction were estimated with the ligand energy inspector function of Molegro Molecular Viewer.

### Bioinformatic analysis

All the protein sequences were retrieved from UniProt with the following entries: Q63633 (*Rattus norvegicus*), A0A076FSX1 (*Mus musculus*), Q9H2X9-2 (*Homo sapiens*), M1EVR1 (*Danio rerio*) and H9GCJ9 (*Anolis carolinensis*). The sequences have been compared on CLUSTAL software.

### Lipid monolayers

The surface pressure of GM1 (Matreya), GM3 (Matreya) or POPC (Biovalley) monolayers was measured with a fully automate microtensiometer (μTROUGH SX, Kibron Inc.) as previously described^70^. Monolayer of the indicated ganglioside or phospholipid was spread on ultrapure apyrogenic water (BioRad) subphases (800 µL) from chloroform/methanol (2:1, vol:vol) or hexane/chloroform/ethanol (11:5:4, vol:vol). After spreading of the film, 5 minutes was allowed for the solvent evaporation. WT-KCC2-GBD or W318S-KCC2-GBD (synthetic peptides with a purity > 95%, Schafer-N) were injected into the subphase and pressure increases produced was continuously recorded. The data were analysed with the FilmWareX 3.57program (Kibron Inc.). The accuracy of the system under our experimental conditions was ±0.25 mN/m for surface pressure.

### Cell Culture

#### HEK293 culture

Human embryonic kidney 293 cells (HEK293T) were obtained from ATCC and maintained in Dubelco Modified Eagle’s Medium (DMEM; ThermoFisher), supplemented with 10% of fetal bovine serum (Gibco), 1 % of L-glutamine (Gibco) and 50 IU.mL^-1^ penicillin-streptomycin (Gibco).

#### Neuro2a culture

Neuro2a cells were obtained from ATCC and were grown in Dubelco Modified Eagle’s Medium (DMEM; ThermoFisher), supplemented with 10% of fetal bovine serum (Gibco), 1 % of L-glutamine (Gibco) and 50 IU.mL^-1^ penicillin-streptomycin (Gibco).

#### Neuronal culture

All animal procedures were carried out according to the European Community Council directive of 24 november 1986 (86/609/EEC), the guidelines of the French Ministry of Agriculture and the Direction Départementale de la Protection des Populations de Paris (Institut du Fer à Moulin, Animalerie des Rongeurs, license C 72-05-22) or the Finnish Ministry (license KEK23-021). All efforts were made to minimize animal suffering and to reduce the number of animals used.

Hippocampal neuron cultures were prepared as previously described^71^ with some modification. Cells were plated at a density of 50,000 cells per well in a 24-well plate on 13 mm round coverslips (Avantor SienceCentral) for imaging previously coated overnight with Poly-L-Lysine Hydrobromide (Sigma-Aldrich) diluted 1:10 in MQ water. Cells were cultured in Neurobasal medium (Gibco) supplemented with 1% Penicillin-Streptomycin (Gibco), 1 % L-Glutamine (Gibco), and 2 % B-27 Supplement (Gibco). The complete Neurobasal medium was pre-equilibrated in the incubator for 2 hours prior to use. Neurons were maintained until day *in vitro* 14 (DIV14) before use, with medium changes twice per week.

All cell cultures were tested for mycoplasma contamination by DAPI, microbiological culture and PCR assays. All cell cultures were maintained at 37 °C and 5 % of CO_2_.

### DNA constructs

The KCC2-Flag construct was generated by insertion of three Flag sequences in the second predicted extracellular loop of KCC2 and was shown to retain normal traffic and function in transfected hippocampal neurons^27^. The WT-KCC2-mCherry, WT-KCC2-pHext, A/A-KCC2-pHext and ΔNTD-KCC2-pHext were gifted by Dr. Igor Medina^48^. Both KCC2-Flag and KCC2-pHext with Tryptophane residue 318 mutated to Serine were generated by GeneCust. Tryptophane nucleotide sequence TGG was changed to Serine nucleotide sequence TCG.

### Transfection and transduction

HEK293 cells were transfected with the appropriate WT-KCC2-mCherry or W318S-KCC2-mCherry plasmid DNAs using Lipofectamine reagent 2000 (Life Technologies) according to manufacturer’s instructions with 1.5 µg of plasmid DNA per 960 mm^2^ well. The transfected cells were used 40 – 44 hours after transfection. For each construct, transfection efficacy was quantified by microscopy.

Neuro2a cells were transfected with WT-KCC2-pHext, or W318S-KCC2-pHext or ΔNTD-KCC2-pHext or T906A/T1006A-KCC2-pHext plasmid DNA using Lipofectamine reagent 2000 (Life Technologies) according to manufacturer’s instructions. The amount of DNA was of

0.75 µg per 24 well plate on 13 mm round coverslips (Avantor SienceCentral) for imaging previously coated overnight with Poly-L-Lysine Hydrobromide (Sigma-Aldrich) diluted 1:10 in MQ water. The transfected cells were used 40 – 44 hours after transfection.

Neuronal cultures were transfected using Lipofectamine 2000 with a maximum of 0.75 µg total plasmid DNA per well, according to manufacturer’s instructions. A plasmid encoding eGFP (0.25 µg) was co-transfected with either a mock plasmid (pCDNA3.1, 0.5 µg), shRNA against KCC2 (0.5 µg) or WT-KCC2-mCherry or W318S-KCC2-mCherry plasmid (0.5 µg). Transfections were carried out in serum-free Neurobasal medium supplemented with 10 mM Mg²⁺. The transfected cells were used until 72 hours after transfection.

### Pharmacology

The following drugs were used: D-threo-PPMP (10 µM; Sigma-Aldrich), dynamin inhibitor (myristoylated peptide, 50 M; Tocris Bioscience), GM1 (10 µM; Matreya). For QD-SPT experiments, coverslips were mounted in a recording chamber. Before imaging, cells were incubated with the PPMP 48 hours at 37 °C in CO_2_ incubator, peptide at 37 °C for 15 min in the imaging medium (see below). They were then imaged for 45 min in the presence of the drugs or peptide. The imaging medium consisted of phenol red-free minimal essential medium supplemented with glucose (33 mM; Sigma) and HEPES (20 mM), glutamine (2mM), Na+-pyruvate (1 mM), and B27 (1×) from Invitrogen.

For immunocytochemistry, drugs were directly added to the culture medium and incubated for 48 hours in a CO_2_ incubator set at 37 °C before cell fixation. The imaging medium was composed of phenol red-free MEM supplemented with glucose (33 mM; Sigma), HEPES (20 mM), glutamine (2 mM), Na+-pyruvate (1 mM), and B27 (1x) (all from Invitrogen).

For electrophysiology, primary neurons were treated with 10 µM PPMP for 48 hours prior to time points in a 5% CO_2_ incubator set at 37 °C.

For live immunostaining, Neuro2a cells were incubated during the transfection, and for 48 hours with 10 µM PPMP.

### Live cell staining for single-particle imaging

Neurons were incubated for 6 min at 37 °C with primary antibodies against Flag (mouse, 1:300, Sigma, cat #F3165), washed, and incubated for 6 min at 37 °C with biotinylated Fab secondary antibodies (goat anti-mouse, 1:300; Jackson Immuno research, cat #115-067-003, West Grove, USA) in imaging medium. After washes, cells were incubated for 1 min with streptavidin-coated quantum dots (QDs) emitting at 655 nm (1 nM; Invitrogen) in PBS (1 M; Invitrogen) supplemented with 10% Casein (v/v) (Sigma).

### Single-particle tracking and analysis

Cells were imaged as previously described^72^ using an Olympus IX71 inverted microscope equipped with a 60X objective (NA 1.42; Olympus) and a 120W Mercury lamp (X-Cite 120Q, Lumen Dynamics). Individual images of GFP-cells, and QD real time recordings (integration time of 30 ms over 1200 consecutive frames) were acquired with an ImagEM EMCCD camera and MetaView software (Meta Imaging 7.7). Cells were imaged within 45 min. QD tracking and trajectory reconstruction were performed with homemade software (Matlab; The Mathworks, Natick, MA) as described in^27,72^. One to two sub-regions of dendrites were quantified per cell. In cases of QD crossing, the trajectories were discarded from analysis. Values of the mean square displacement (MSD) plot vs. time were calculated for each trajectory by applying the relation:

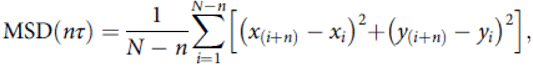

where τ is the acquisition time, N is the total number of frames, n and i are positive integers with n determining the time increment. Diffusion coefficients (D) were calculated by fitting the first four points without origin of the MSD vs. time curves with the equation: MSDðnτÞ ¼ 4Dnτ þ b; where b is a constant reflecting the spot localization accuracy. Depending on the type of lamp used for imaging, the QD pointing accuracy is ∼20–30 nm, a value well below the measured explored areas (at least 1 log difference). Synaptic dwell time was defined as the duration of detection of QDs at synapses on a recording divided by the number of exits as detailed previously^73^. The explored area of each trajectory was defined as the MSD value of the trajectory at two different time intervals of at 0.42 and 0.45 s.

### Immunocytochemistry

#### For KCC2 clusters study at the membrane

WT-KCC2-Flag-ires-GFP or W318S-KCC2-Flag-ires-GFP membrane expression and clustering were assessed with staining performed after a 4 min fixation at room temperature (RT) in paraformaldehyde (PFA; 4% w/v; Sigma) and sucrose (20% w/v; Sigma) solution in PBS. Cells were washed three times in PBS and incubated for 30 min at RT in a blocking solution composed of goat serum (GS; 10% v/v; Invitrogen) prepared in PBS. Neurons were then incubated for 60–180 min at RT with Flag (mouse, 1:400, Sigma) in PBS–GS (GS; 3% v/v; Invitrogen) blocking solution. After washing, neurons were incubated with Cy™3 AffiniPure Donkey Anti-Rabbit IgG (H + L) (1.9 g/mL; Jackson ImmunoResearch) for standard epifluorescence assays, The coverslips were then washed and mounted on slides with mowiol 844 (48 mg/mL; Sigma). Sets of neurons compared for quantification were labeled and imaged simultaneously.

#### For live immunostaining of Neuro2a

Cells were stained as previously described^48^ with minor adjustments. Cells were incubated with polyclonal chicken anti-GFP antibody (1:500; Aves GFP-1010) for 2 hours at 37 °C. After labeling, cultures were transferred at RT to HEPES-buffered saline and placed for 10 min into the thermo-isolated box at 13 °C. The cells were then incubated at 13 °C for 20 min with a goat anti-chicken Alexa Fluor Plus 555-conjugated antibody (1:500, Invitrogen A-32932). After rinsing in a HEPES-buffered saline solution for 10 min at 13°C, the cells were fixed in 4 % PFA for 20 min at RT, permeabilized (in 0.3 % Triton X-100 solution), blocked (5% goat serum solution) for 30 min, and labeled with a second secondary antibody, goat anti-chicken Alexa Fluor 647 (Invitrogen A-21449) for 1 hour. To detect the total level of ectopic KCC2-related proteins expressed, the cells were labeled with mouse anti-GFP antibody (Novus NB600-597SS) for 1 hour at RT) and revealed with anti-mouse Alexa 488 antibody (Invitrogen A-11001) for 1 hour at RT.

#### For KCC2, GM1 and cholesterol colocalization staining

Live neurons were incubated with an anti-GM1 antibody (mouse, 1:250, Cayman) and a homemade cholesterol probe (His₆-mCherry-tagged perfringolysin theta toxin D4^40^, 1:250) for 2 hours at 37 °C. Neurons were then fixed with PFA 4% for 5 minutes, washed twice with PBS and blocked in a staining buffer (SB) containing 2.5% BSA, 2.5% NHS, and 5% glycine in 1x PBS. Neurons were incubated for 2 hours at RT with a Cy5-conjugated Donkey Anti-Mouse antibody (1:500, Biosite 715-175-150). After washing, neurons permeabilized in SB containing 0.5% Triton X-100 for 30 minutes. KCC2 was labeled using a rabbit anti-KCC2 antibody^36^ (1:1000) in SB + 0.3% Triton X-100 overnight at 4 °C. After washing, neurons were incubated with an Alexa Fluor Plus 488 Donkey Anti-Rabbit (1:500, Invitrogen A-32790) for 2 hours at RT. After washing, neurons were incubated with a Rat anti-RFP antibody (1:1000, ChromoTek 5f8) overnight at 4 °C in SB + 0.3% Triton X-100. On the last day neurons were washed and incubated with an Alexa Fluor 555 Goat Anti-Rat antibody (1:500, Invitrogen A-21434) for 2 hours at RT. The coverslips were then washed and nuclei were stained with DAPI (1:500, Roche Life Science) for 30 minutes. Finally, neurons were washed and mounted using Fluoromount-G™ Mounting Medium (Invitrogen).

### Fluorescence Image Acquisition and Analysis

#### For KCC2 clusters study at the membrane

Image acquisition was performed using a 100 objective (1.40 NA) on a Leica (Nussloch, Germany) DM6000 upright microscope with a 12-bit cooled CCD camera (Micromax, Roper Scientific) using MetaMorph software Series 7.8 (Roper Scientific). To assess KCC2-Flag clusters, the exposure time was determined on the brightest experimental condition in order to be non-saturating and was fixed for all cells and conditions to be analyzed. Quantification was performed using a MetaMorph routine (Roper Scientific). For the dendritic intensity and clustering analysis, a region of interest (ROI) was traced around a selected dendrite, and the average pixel intensity in the ROI was measured. For the clustering analysis, images were filtered using the flattened background (kernel size,3 x 3 x 2) function on Metamorph to enhance cluster outlines, and an intensity threshold defined by the user was set to identify the clusters and avoid their coalescence. Clusters were outlined, and the corresponding regions were transferred onto the raw images to determine KCC2-Flag cluster number, surface, and fluorescence intensity. The area of the dendritic region analyzed was used to calculate the number of clusters per surface area. We analyzed ∼15 cells per experimental condition and per culture.

#### For live Imaging of Neuro2a cells

Images were acquired with an Olympus Fluorview-500 confocal microscope using oil-immersion objective 40× Plan-Apochromat (NA 1.4) Oil DIC (UV) Vis-IR M27, zoom 3, 26nm in X; Y resolution and 500nm in Z resolution. The bit depth used was 8bit. We randomly selected and focused on a transfected cell by visualizing only Alexa Fluor 488 fluorescence and then acquired z-stack images of Alexa Fluor 488, Alexa Fluor 555, and Alexa Fluor 647 fluorochromes emitted fluorescence using, respectively, green (excitation 493 nm, emission 400–550 nm), red (excitation 553 nm, emission 535–617 nm), and infrared (excitation 653, emission 645-700 nm) channels of the microscope. Each z-stack included 15-25 planes of 0.5-µm optical thickness and 0.5 µm distance between planes. The fluorescence intensities of each cell were analyzed with ImageJ 1.54p. First, a Z-projection (SUM) was generated for all channels. To isolate the Alexa Fluor 647 signal not overlapping with Alexa Fluor 555 fluorescence, an image subtraction was performed between the Alexa Fluor 647 and Alexa Fluor 555 channels. This resulted in the "internalized pool" images. Next, regions of interest (ROIs) were defined for each cell based on the Alexa Fluor 488 channel to ensure the analysis was restricted to transfected cells. The mean fluorescence intensity was then measured for each channel within the ROIs. The same analysis parameters were consistently applied across all experiments.

#### For KCC2, GM1 and cholesterol colocalization

Image acquisition was performed on an LSM 980 Axio Examiner confocal microscope using a 63x oil immersion objective (Objective alpha Plan-Apochromat 63x/1.46 Oil Corr M27). Pixel resolution was set to 82 nm in the x and y dimensions, and 120 nm in z. A zoom factor of 1.5x was used for each neuron. To optimize colocalization, the pinhole for each channel was adjusted to obtain a 0.4 µm optical section. The pixel time was 1.91 µs (2x averaged per line). The images were saved in 16bit depth with a resolution of 1101×1101 pixels. A bidirectional scan method was used to reduce the time of acquisition of each picture. The detector type used were Gallium arsenide phosphide cathodes-photomultiplier tubes (GaAsp-PTM).

The raw data were loaded on Huygens Professional 24.10.0p6 64b, cells ROIs were extracted from the rest of the images by using KCC2’s signal with the following setting, the thresholds between 1 to 5% and the seed at 100%. All channels were first deconvoluted using the deconvolution express in standard mode. Picture were cropped to keep only the target branch and were deconvoluted a second time in aggressive mode. For each neuron, 1 to 2 primary branches were quantified. For each channel, the total number of voxels is detected using the “Watershed” mode with the following settings: 1% threshold, seed at 0% and fragmentation at 100%. To obtain the ratio between the total voxel and the voxels involved in the different double colocalization possibilities, colocalization images were created using the analysis tool (Roi-Intersection-Remove outside), all the pixels that are colocalizing were kept with them respective intensities in every channel. Two pictures were made as results, the pixels colocalizing between GM1-Cholesterol and GM1-KCC2. Then after creating a composite picture of the two previous, the colocalization procedure was repeated, resulting only in the pixel triple colocalizing between GM1-Cholesterol-KCC2.

### Brain samples

Wistar young rats were housed are maintained in a 12 h light/12 h dark cycle environment with controlled temperature (23 ± 2°C), food and water were given *ad libitum*. Animals were deeply anaesthetized with isoflurane and then injected with 7 % chloralhydrate (100 µL per 100 grams). Brains were collected and hemispheres were frozen in liquid nitrogen and stored at -80 °C. All experimental procedures were in accordance with-and have been validated by a French ethical committee.

For KCC2, flotillin-1 and transferrin-receptor expression quantification, C57Bl6 St3Gal5 mice (8 weeks old) were mate and maintained at Neuroscience Center, HiLIFE animal facility under controlled environment (23 ± 2°C; 12 hours light/dark cycle and ad libitum access to food and water). All experimental procedures were validated beforehand by local ethical committee (decision number: KEK23-016).

### Genotyping

St3Gal5 mice were genotyped from ear samples with AccuStart II Mouse Genotyping Kit (Quantabio). Samples in 50 µL of extraction reagent were incubated at 95 °C for 30 min. After cooling at RT, 50 µL of stabilization buffer was added. PCR was done with 1:10 dilution in sterile water (Invitrogen). The PCR mix was composed by 1:80 of primers and 1:2 of Accustart 2X supermix, completed by sterile water. Two PCR were done in parallel: 1 wild-type (WT) PCR and 1 mutant PCR. Primers for WT PCR were 5’-AGC TCA GAG CTA TGC TCA GGA-3’ and 5’-TAC CAC ATC GAA CTG GTT GAG-3’. Primers for mutant PCR were 5’-CAA TAG ATC TGA CCC CTA TGC-3’ and 5’-TCG CCT TCT TGA CGA GTT CTT CTG AG-3’. Diluted samples were added in 1:20 to the mix. The WT PCR program was: 1) 95°C 3 min 2) 94°C 30 sec 3) 58°C 30 sec 4) 72°C 30 sec 5) repeat steps 2-4 39 times for a total of 40 cycles 6) 72°C 10 minutes. The mutant PCR program was: 1) 95°C 3 min 2) 94°C 20 sec 3) 51°C 25 sec 4) 72°C 30 sec 5) repeat steps 2-4 39 times for a total of 40 cycles 6) 72°C 10 minutes. The PCR products were run in a 2% agarose gel (Nippon Genetics) in TAE 1X pH 8,0 with 1:25 of Midori Green Xtra (Nippon Genetics) at 100 V for 30 min. The WT band was seen at 337 bp and the mutant band at 250 bp.

### Sucrose gradient

HEK-293T cells were transfected with either WT-KCC2-mCherry or W318S-KCC2-mCherry constructs and subsequently treated with PPMP (10 µM) for 48 h or methyl-β-cyclodextrin (1 mM) for 24 h. Following incubation, cells were washed twice with PBS, harvested in PBS, and centrifuged at 2,000 × g for 5 min at room temperature. Cell pellets were lysed in 500 µL of TKM buffer (0.5 mM Tris, 25 mM KCl, 1 mM EDTA) supplemented with protease and phosphatase inhibitors. An aliquot corresponding to one-tenth of the lysate was retained as the total fraction. A total of 2 mg of protein was applied to a discontinuous sucrose gradient (80%, 40% [containing the lysate], 36%, and 5%) and centrifuged for 18 h at 171,100 × g. Fractions (1 mL) were collected and precipitated overnight at −20 °C using cold acetone (1:1, acetone:sample). Samples were then centrifuged at 14,000 × g for 10 min at 4 °C. Resulting pellets were resuspended in 30 µL of IP lysis buffer and processed for SDS-PAGE and immunoblot analysis.

### Immunoprecipitation and immunoblotting

All biochemical preparations and centrifugation were performed at 4 °C. Rat brains (PND5, PND15 and PND30), mice brains and transfected HEK293 cells were homogenized in cold IP Buffer (Tris-HCl, 50 mM; pH8; NaCl, 100 mM; <u>NP40 1%</u>; MgCl_2_, 1 mM; protease inhibitor EDTA free) using a homogenizer. Samples (400 - 500 µg of proteins) were incubated for 2 hours in IP Buffer with the dedicated antibody (against GM1 or KCC2 or no antibody) and with agitation. Then solutions were incubated overnight with 60 µL protein-A-Sepharose (Sigma). Following antibody-antigen binding to the beads, the beads were washed three times with IP Buffer 100 (NaCl, 100 mM), twice with IP Buffer 500 (NaCl, 500 mM) and twice with IP Buffer 100 (NaCl, 100 mM), then stocked in PBS.

One fraction of bound proteins (IP fractions) was eluted with SDS sample buffer containing DTT and then separated in NuPAGE 4-12% Bis-Tris gels (Invitrogen). Gel was equilibrated by washing wells with MQ water and pre-run for 1 hour at 200 V at 4 °C in Running Buffer Bolt MES SDS 1X (Invitrogen). Then, samples were run at 4 °C at 100 V for 15 min then 200 V for 1h30. After the run, wet transfer was done on a PVDF membrane (0,45 µm) from GE HealthCare, for 2 hours at 140 V at 4 °C. Membrane was rinsed in TBST (TBS 1X pH 7,4 + Tween 20 0,1 % (Sigma-Aldrich)) for 5 min. The membrane was blocked for 1 hour at RT in saturation buffer (BSA 5 % (biowest) in TBST) with agitation. Then primary antibodies were incubated overnight at 4 °C with agitation: antibody anti-KCC2 (homemade) at 1:3000 and antibody anti-GAPDH (Abcam) at 1:20000 in 2,5 % BSA in TBST. Then, membrane was washed 3 times for 15 min in TBST with agitation and secondary antibodies were incubated 1 hour at RT with agitation: anti-rabbit (Cytiva) at 1:10000, anti-mouse (Cytiva) at 1:1000 in 2,5 % BSA in TBST. Then membrane was washed 3 times for 15 min in TBST with agitation and membrane was detected by Enhanced Chemiluminescence (Thermo Scientific) in Syngene G:BOX – F3 (Syngene) with GENESys software (version 1.8.11.0). Analysis was performed using Gel Plot Analyzer plugin (ImageJ).

For the rat brain IP fractions, the second fraction of bound proteins was centrifugated and the pellet was treated with Folch solvent (chlorofom/methanol, 2/1, v/v). The lipid fractions were deposited on HPTLC silica plate and treated with isobutylmetacrylate(Sigma). After saturation with gelatin-A (Sigma), plates were incubated with anti-GM1 antibody (1:650) for 2 hours at RT and with secondary Alexa 488 antibody (1:500) for 1 hour at RT. For both lipids and proteins, signals were revealed on Syngene G:BOX – F3 (Syngene) with GENESys software (version 1.8.11.0).

### Efficacy of KCC2-mediated Cl^−^ extrusion

#### Whole cell patch-clamp

Somatic recordings in hippocampal neurons (DIV 13–15) were performed in standard extracellular solution at 34 °C in the whole-cell voltage-clamp configuration using a patch-clamp amplifier, EPC 10 patch-clamp amplifier (HEKA Elektronik, Inc., Germany). The composition of the extracellular solution was (in mM): 127 NaCl; 3 KCl; 2 CaCl2; 1.3 MgCl2; 10 D-glucose; 20 HEPES; pH 7.4 was adjusted with NaOH solution. Patch pipettes were fabricated from borosilicate glass (Harward Apparatus, UK), and their resistances ranged from 4 to 8 MOhm. Composition of the patch pipette solution was (in mM): 18 KCl; 111 K-gluconate; 0.5 CaCl2; 2 NaOH; 10 glucose; 10 HEPES; 2 Mg-ATP; 5 BAPTA; pH 7.3 was adjusted with KOH solution. Membrane potential was clamped at -70 mV. All membrane potential values were corrected for a liquid-junction potential of 14.16 mV. Before recording, the first 5 minutes were dedicated to load the whole cell with the intracellular solution. The reversal potential of GABAergic currents (E_GABA_) was determined from the current–voltage (I–V) relation obtained by using ramp voltage protocol. NKCC1 was blocked throughout the experiments with 10 μM bumetanide, action potentials with 1 μM TTX and GABA_B_ receptors with 1 μM CGP 35348. ***Ionophoresis.*** For local iontophoretic application of GABA, brief (100 ms) positive current pulses (50–70 nA) were delivered by a sharp micropipette (20-25 MOhm when filled with 250 mM GABA in 250 mM HCl). Iontophoretic GABA injections were given not more often than once in 1 min. Constant negative current of −5 nA was applied to the micropipette to compensate for the passive leak of GABA. GABA was applied at the soma and at the dendrite (∼100 μm from the soma) of a given neuron.

### Data and statistical analysis

All the statistical analyses were performed using GraphPad Prism (version 10) software. Methodology schematics was generated using BioRender (Biorender.com).

## Supporting information

Supplementary informations

## Author contributions

C.K., A.P.dl.C, A.A., E.B., O.A., A.T., Se.L., J.A., F.M., M.R. and C.D.S. performed experiments. C.K., A.P.dl.C, A.A., E.B., O.A, A.T., J.A., M.R., Sa.L., C.R and C.D.S. analyzed and interpreted the data. C.D.S. and C.R. designed the study Sa.L., C.R. and C.D.S. secured funding. C.D.S wrote the paper. All the authors have read and accepted the present manuscript.

## Acknowledgment

We thank the INMED platforms, especially Dr. François Michel from INMAgic for his help with confocal microscope and image analysis; the PBMC platform for providing the equipment’s for immunoblotting and cell cultures and the neuronal cell culture unit from the Neuroscience Center, HiLIFE. We thank Marie Kruz for excellent technical support with the animals and the animal facility from Neuroscience Center HiLIFE. We thank Dr. Igor Medina for giving us the constructs necessary to this study. The mouse strain used for this research project, B6;129S-St3gal5tm1Rlp/Mmmh, RRID:MMRRC_000374-MU, was obtained from the Mutant Mouse Resource and Research Centers (MMRRC) at the University of Missouri, an NIH-funded strain repository, and was submitted to the MMRRC by Richard L. Proia, Ph.D., National Institutes of Health, National Institute of Diabetes and Digestive and Kidney Diseases.

This work was supported by INSERM, CNRS, the Agence Nationale de la Recherche ANR-GABGANG to C.R., Sa.L and C.D.S, by the Research Council of Finland to C.R (decision 341361) and to C.D.S (decisions 333096, 335956 and 358122) and by the Fondation pour la Recherche Médicale EQU202203014844 to Sa.L.. Equipment in Sa.L. laboratory was supported by DIM C-Brains from Région Ile-de-France and by the Fondation pour la Recherche sur le Cerveau / Rotary ‘Espoir en tête’. The team *The dynamic Synapse* is affiliated with PSL-NEURO, funded by PSL University. This work was also supported by grants from the *‘Gueules Cassées’* foundation, *‘Ligue Française Contre l’Epilepsie’*, *‘Fondation Française pour la Recherche sur l’Epilepsie’* and the Sigrid Jusélius Foundation to C.D.S.. Epilepsiatutkimussäätiö supported C.K., A.P.dl.C., and E.B, and, Instrumentarium foundation C.K.. A.P.dl.C. also received support from the EDUFI. We also want to thank the Neuroscience Center and the HiLIFE fellow program for supporting C.D.S. as a young group leader.

