## Supplementary informations for "Ganglioside GM1-enriched rafts regulate the neuronal chloride co-transporter KCC2"

### Material and methods

**Lipid monolayers.** The surface pressure of GM1 (Matreya), GD1a (Matreya), GD1b (Matreya) or GT1b (Matreya) monolayers was measured with a fully automate microtensiometer (µTROUGH SX, Kibron Inc.) as previously described[1]. Monolayer of the indicated ganglioside was spread on ultrapure apyrogenic water (BioRad) subphases (800 µL) from chloroform/methanol (2:1, vol:vol). After spreading of the film, 5 minutes was allowed for the solvent evaporation. WT-KCC2-GBD (synthetic peptide with a purity > 95%, Schafer-N) were injected into the subphase and pressure increases produced was continuously recorded. The data were analysed with the FilmWareX 3.57program (Kibron Inc.). The accuracy of the system under our experimental conditions was ±0.25 mN/m for surface pressure.

**Animals.** For KCC2, GM1 and cholesterol quantification, 1 month-old male Wistar rats were purchased from Charles River (Lyon, France) and maintained at INMED animal house facilities under controlled environment (23 ± 2°C; 12 hours light/dark cycle and ad libitum access to food and water).

For KCC2, flotillin-1 and transferrin-receptor expression quantification, C57Bl6 St3Gal5 mice (8 weeks old) were mate and maintained at Neuroscience Center, HiLIFE animal facility under controlled environment (23 ± 2°C; 12 hours light/dark cycle and ad libitum access to food and water).

Rats and mice were handled according to European Union guidelines (2010/63) after approval by the Animal's Ethics Committee. All efforts were made to reduce the number of animals used and to minimize their stress and discomfort, following ARRIVE guidelines.

**Cell Culture.** ***HEK293T culture.*** Human embryonic kidney 293 cells (HEK293T cells) were obtained from ATCC and maintained in Dubelco Modified Eagle's Medium (DMEM; ThermoFisher), supplemented with 10% of fetal bovine serum (Gibco), 1 % of L-glutamine (Gibco) and 50 $IU.mL^{-1}$ penicillin-streptomycin (Gibco).

***Neuro2a culture.*** Neuro2a cells were obtained from ATCC and were grown in Dubelco Modified Eagle's Medium (DMEM; ThermoFisher), supplemented with 10% of fetal bovine serum (Gibco), 1 % of L-glutamine (Gibco) and 50 $IU.mL^{-1}$ penicillin-streptomycin (Gibco).

***Neuronal culture.*** All animal procedures were carried out according to the European Community Council directive of 24 november 1986 (86/609/EEC), the guidelines of the French Ministry of Agriculture and Finnish ethical committee. All efforts were made to minimize animal suffering and to reduce the number of animals used.

Hippocampal neuron cultures were prepared as previously described[2] with some modification. Cells were plated at a density of 50,000 cells per well in a 24-well plate on 13 mm round coverslips (Avantor SienceCentral) for imaging previously coated overnight with Poly-L-Lysine Hydrobromide (Sigma-Aldrich) diluted 1:10 in MQ water. Cells were cultured in Neurobasal medium (Gibco) supplemented with 1% Penicillin-Streptomycin (Gibco), 1 % L-Glutamine (Gibco), and 2 % B-27 Supplement (Gibco). The complete Neurobasal medium was pre-equilibrated in the incubator for 2 hours prior to use. Neurons were maintained until day *in vitro* 14 (DIV14) before use, with medium changes twice per week.

All cell cultures were tested for mycoplasma contamination by DAPI, microbiological culture and PCR assays. All cell cultures were maintained at 37 °C and 5 % of $CO_2$.

**Cell mitochondrial activity.** Mitochondrial activity was assessed using 3-(4,5-dimethylth-iazol-2-yl)-5-(3-carboxymethoxyphenyl)-2-(4-sulfophenyl)-2H- tetrazolium (MTS) assay (Promega). Cells were exposed to various concentrations of PPMP (from 0 to 100 µM for 48 h). Then MTS was added (20 mL/100 mL) and the cells were incubated for 3 h at 37 °C. MTS was spectrophotometrically measured at 490 nm. Survival of the control cells not exposed to PPMP was set at 100% and treated groups were represented as percentage of control values.

**Transfections.** HEK293T cells were transfected with the appropriate WT-KCC2-mCherry or W318S-KCC2-mCherry plasmid DNAs using Lipofectamine reagent 2000 (Life Technologies) according to manufacturer's instructions with 1.5 µg of plasmid DNA per 960 $mm^2$ well. The transfected cells were used 40 – 44 hours after transfection. For each construct, transfection efficacy was quantified by microscopy. For each picture, total cells and number of cells expressing mCherry were counted thanks to ImageJ Cell counter plugging to determine ratio transfection.

Neuro2a cells were transfected with WT-KCC2-pHext, or W318S-KCC2-pHext or ΔNTD-KCC2-pHext or T906A/T1006A-KCC2-pHext plasmid DNA using Lipofectamine reagent 2000 (Life Technologies) according to manufacturer's instructions. The amount of DNA was of 0.75 µg per 24 well plate on 13 mm round coverslips (Avantor SienceCentral) for imaging previously coated overnight with Poly-L-Lysine Hydrobromide (Sigma-Aldrich) diluted 1:10 in MQ water. The transfected cells were used 40 – 44 hours after transfection.

**Immunocytochemistry.** ***Immunostaining of primary cultures.*** Primary neuronal cultures at various days of division were fixed with PFA 4% for 1 hour at 4 °C. Then saturation was done by incubating the cells with a solution of 2.5% BSA and 2.5% NGS for 1 hour at RT. The cells were incubated with anti-KCC2 antibody (1:5000; home-made[3]) over night at 4 °C, washed and incubated with Donkey Anti-Rabbit Alexa 555 antibody (1:500; Invitrogen A31572) 2 hours at RT, fixed 20 minutes with cold PFA 4%. After a new saturation step of 2 hours at RT with a solution of 2.5% BSA and 2.5% NGS, cells were incubated with anti-GM1 antibody (1:650, Matreya) over night at 4 °C. After treatment with Goat Anti-Rabbit Alexa 488 antibody (1:500; Invitrogen A11034), cells were incubated with anti-MAP2 antibody (1:500) or anti-GFAP antibody (1:500) 4 hours at RT. Cytoskeleton staining was then revealed by using the appropriate secondary antibody Alexa 647 (1:500). The nuclei were stained by using Hoechst solution (Sigma). Cells were then visualized tanks to a confocal microscopy (Zeiss; LSM 800) with x40 objective. Pictures were analyzed with ImageJ and JACoP plugin was used to quantify the Pearson's coefficient and the amount of GM1 overlapping KCC2 (M1 Mender's coefficient) or the amount of KCC2 overlapping GM1 (M2 Mender's coefficient)[4].

***GM1 labelling of Neuro2a cells.*** Neuro2a cells were treated or not with 10 µM of PPMP for 48 hours and fixed with cold PFA 4% for 1 hour. After saturation with a solution of 2.5% BSA and 2.5% NGS for 1 hour, cells were incubated with anti-GM1 antibody (Matreya) overnight at 4 °C. Cells were then incubated with Goat Anti-Rabbit Alexa 488 antibody (1:500; ThermoFisher) for 2 hours at RT. The nuclei were stained with Dapi. Cells were observed under a confocal microscope (Zeiss; LSM 800) at the x40 objective. Pictures were analyzed with ImageJ software.

**Lipids and proteins extraction and analysis from rat brains.** GM1, cholesterol and KCC2 expressions were analyzed on rat brain at PND5, PND15 and PND30. For each condition, one hemisphere was dedicated to lipids analysis, and the other one was to proteins analysis. Lipids from rat brains were extracted and purified as previously described[5]. Briefly, lipids were extracted, gangliosides were recovered from the upper phase of a Folch partition extraction and cholesterol from the lower phase. Gangliosides were purified through an exclusion chromatography (Sephadex G25). Lipids were analyzed with high performance thin layer chromatography (HPTLC) and colored with orcinol.

For the protein fraction, hemispheres were homogenized in cold RIPA Buffer using a homogenizer 25 µg of proteins were incubated with SDS, DTT and Laemmli loading Buffer. The samples were separated into 4 -12 % SDS-PAGE gel (Criterion gel, Bio-Rad) and transferred to nitrocellulose membrane (Whatman). After blocking in Tris-buffered saline/0.1% tween/5% BSA, membranes were exposed overnight at 4 °C to primary antibody diluted in blocking solution (Tris-buffered saline/0.1% tween/2.5% BSA), anti-KCC2 (1:5000). Secondary antibody (goat anti-rabbit Alexa 488; 1:500; A11034) was applied for 2 hours at room temperature. For normalization, membranes were incubated with anti-β-tubulin (TUBB3 18020, Biolegend) and secondary antibody (anti-mouse HRP; 1:500; ThermoFisher). For both lipids and proteins, signals were revealed on the image analysis software G Box (Syngene) and quantifications were performed using Gel Plot Analyzer plugin (ImageJ).

**Co-immunoprecipitation.** Experiments were done on ice. HEK293 cells transfected with the appropriate KCC2-construct were lysate with IP buffer 100 mM: Tris HCl 5 mM, NaCl 100 mM, $Mg^{2+}$ 1 mM, Nonidet P40 1% (9016-45-9, Sigma-Aldrich) and protease inhibitor

(A32955, Thermo Scientific), pH 8,0). For each condition, proteins were quantified using a Pierce BCA protein quantification kit (23225, Thermo Scientific). For each construct, 3 tubes were done: α (condition of interest), β (positive control) and γ (negative control). Samples were prepared with 400 µg of proteins diluted in volume total of 75 µL of IP buffer 100 mM We added 1:38,5 of antibody anti-GM1 (32289, Cayman Chemical) in α tube and 1:38,5 of antibody anti-KCC2 (homemade antibody[3]) in β tube (positive control). No antibody was added in γ tube for negative control. Then, tubes were incubated for 2 hours at 4 °C with parafilm and under gentle agitation. After incubation, 78 mg / mL of Protein A-Sepharose Beads (P3391-1.5G, Millipore) were added in all tubes and tubes were incubated overnight at 4 °C with parafilm and under gentle agitation. Approximatively 18 hours after the beginning of the first part of the experiment, samples were centrifuged at 13 00 rpm at 4 °C for 5 min. Then 3 washes were done with IP buffer 100 mM – 150 µL of buffer, pellet is resuspended and centrifugation at 13 000 rpm at 4 °C for 5 min. Then 2 washes were done with 150 µL of IP buffer 500 mM (Tris HCl 5 mM, NaCl 500 mM, $Mg^{2+}$ 1 mM, Nonidet P40 1% and protease inhibitor, pH 8,0), 1 wash with 150 µL with IP buffer 100 mM and 1 wash with 150 µL with PBS -/-. Pellets were stored in PBS -/- at -20°C.

For electrophoresis, proteins were incubated with SDS, DTT and Laemmli loading Buffer. The samples were separated in 4 -12 % SDS-PAGE gel (Criterion gel, Bio-Rad) and transferred to nitrocellulose membrane (Whatman). After blocking in Tris-buffered saline/0.1% tween/5% BSA, membranes were exposed overnight at 4 °C to primary antibody diluted in blocking solution (Tris-buffered saline/0.1% tween/2.5% BSA), anti-KCC2 (1:5000). Secondary antibody (anti-rabbit HRP; 1:500; ThermoFisher) was applied for 2 hours at room temperature. Signals were revealed on the image analysis software G Box (Syngene) and quantifications were performed using Gel Plot Analyzer plugin (ImageJ).

**Data and statistical analysis.** All the statistical analyses were performed using GraphPad Prism (version 10) software.

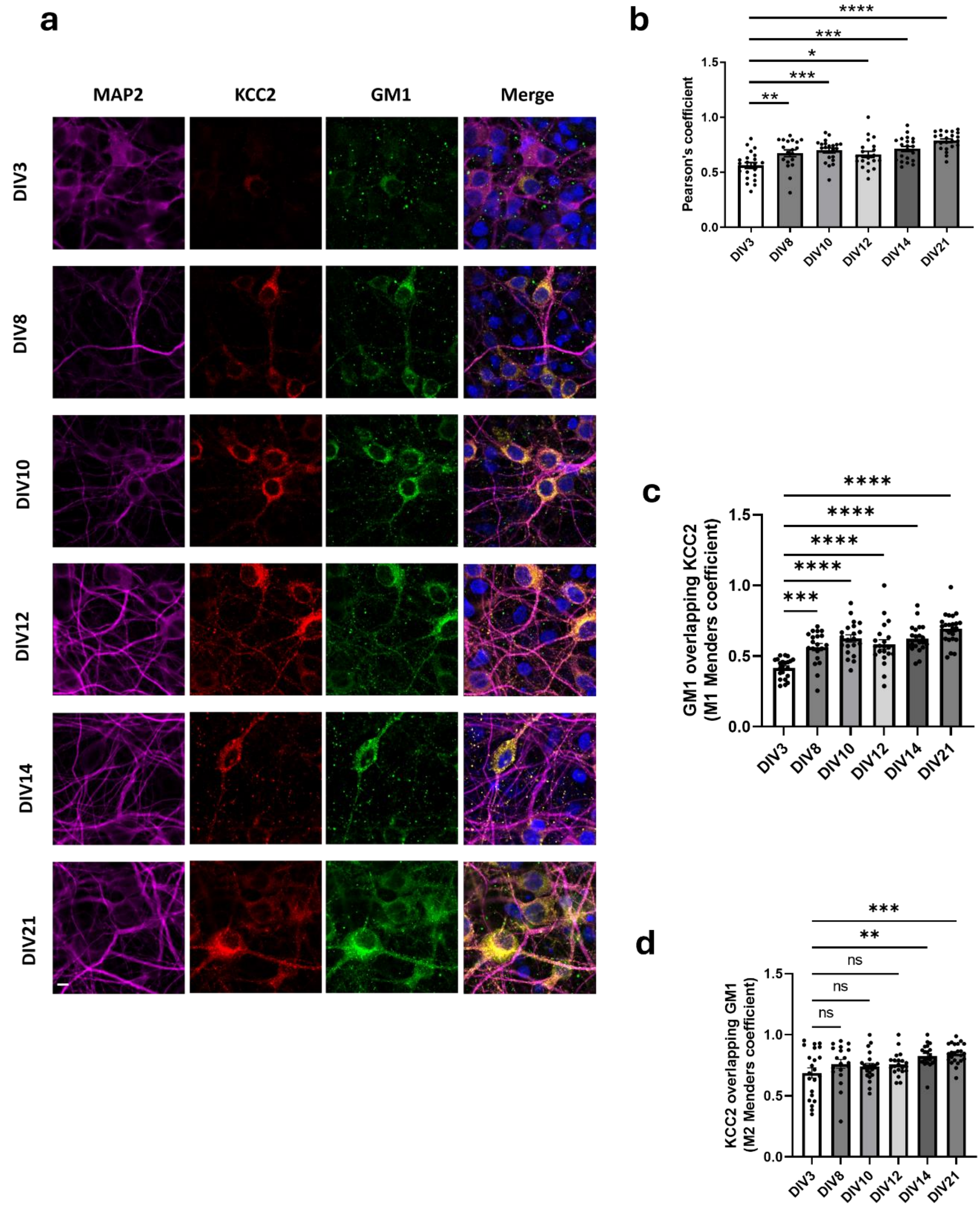


**Figure S1: Spatial-temporal expression of KCC2 and GM1 in hippocampal cultures.**

**a.** Immunostaining of MAP2 (magenta), KCC2 (red) and GM1 (green) in primary hippocampal cultures from 3 to 21 DIV-old. Pictures are representative of three independent experiments made in duplicate. Scale bar: 5 µm.

**b-d** Quantification of the Pearson's coefficient (**b**), GM1 overlapping KCC2 coefficient (**c**), and, KCC2 overlapping GM1 coefficient (**d**). In all graphs, histograms represent means ± SEM of the measured variable (one-way ANOVA; ns – not significant; *p < 0.05; **p < 0.01; ***p < 0.001; ****p < 0.0001).

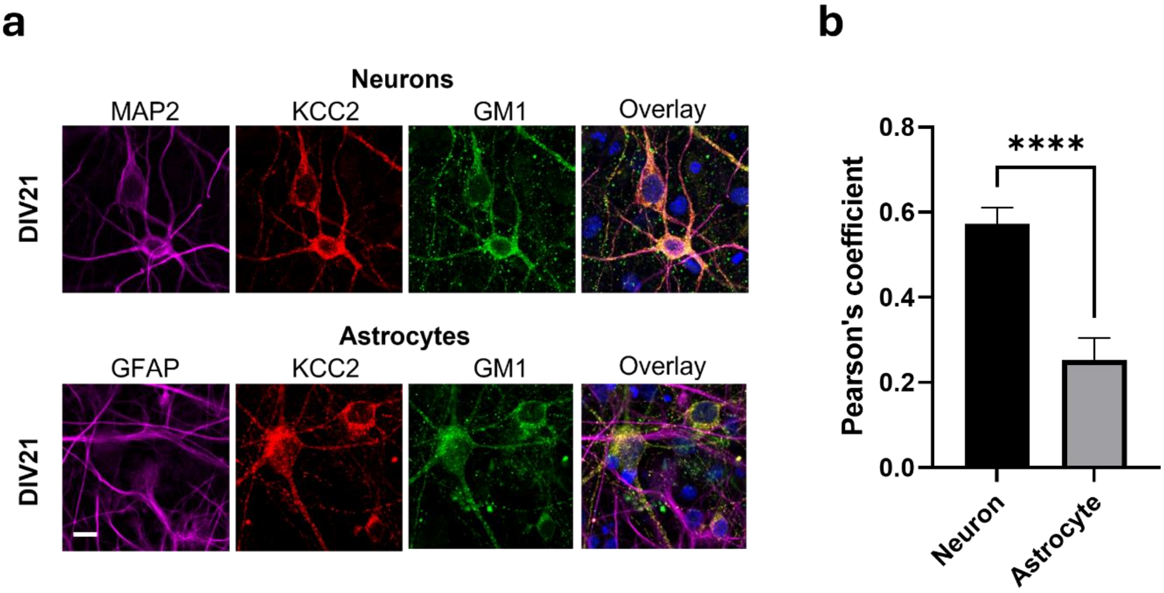


**Figure S2: KCC2 and GM1 colocalize in neurons but not in astrocytes.**

**a.** Immunostaining of MAP2 (magenta), GFAP (magenta), KCC2 (red) and GM1 (green) in primary hippocampal cultures from at 21 DIV-old. Pictures are representative of three independent experiments made in duplicate. Scale bar: 10 µm.

**b.** Quantification of the Pearson's coefficient for KCC2 and GM1. In both graphs, histograms represent means ± SEM of the measured variable, (neurons, n = 45 cells; astrocytes, n = 33 cells), Mann-Whitney test ****$p < 0.0001$.

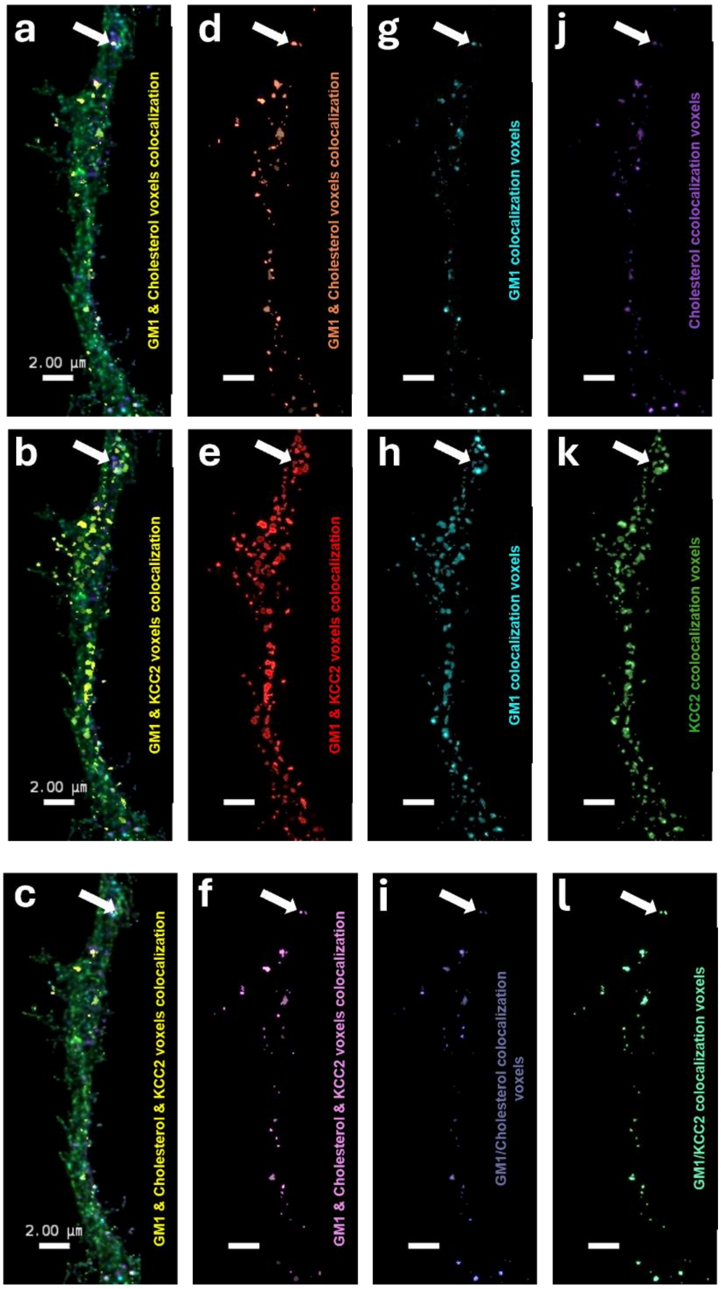


**Figure S3: KCC2 is enriched in cholesterol and GM1 lipid rafts.**

Representative confocal images of primary hippocampal neuron cultures at DIV14, stained with cholesterol probe and antibodies against GM1 and KCC2. **a-c** Colocalization (GM1 & cholesterol; GM1 & KCC2 or GM1 & cholesterol & KCC2) labeled in yellow. **d-f** Artificial channel showing the colocalized pixel in the different configuration. **g-i,** and **j-l** Colocalized pixels in each respective channel obtained by using the artificial channels as a mask to exclude the none involved pixels. Pictures are representative from three independent experiments made in duplicate, between 7 and 8 neurons were analyzed for each batch; for each neuron, 1–2 primary branches were quantified.

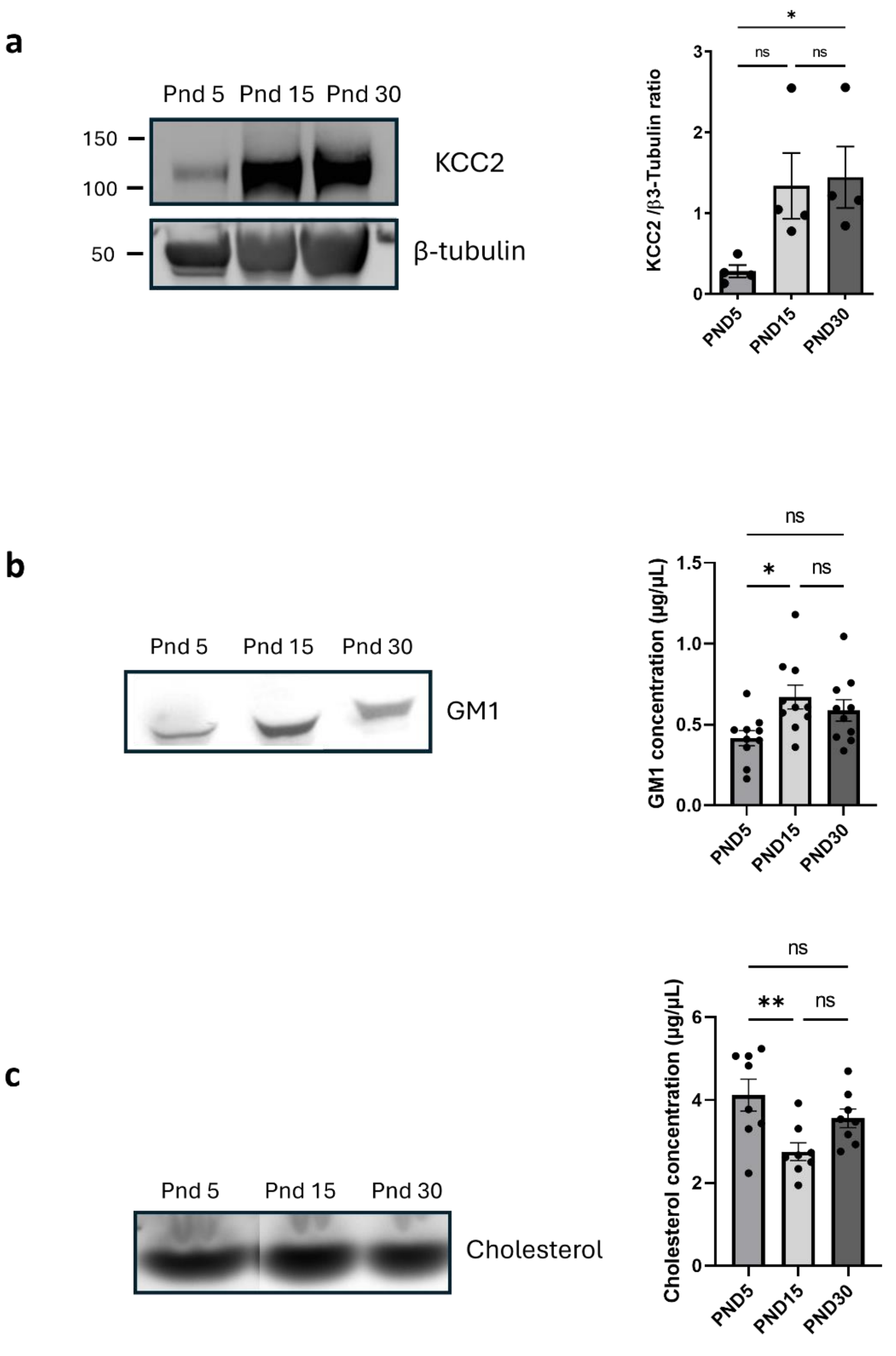


**Figure S4: Developmental expression profile of KCC2 and lipid rafts in rat brains.**

**a.** Western blot and quantification of KCC2 expression in rat brain at PND5, PND15 and PND30. n = 4, histograms represent means ± SEM of the measured variable (Kruskal-Wallis; ns – not significant; *p < 0.05).

**b**. Thin layer chromatography and quantification of GM1 in rat brain at PND5, PND15 and PND30. n = 10, histograms represent means ± SEM of the measured variable (One-way ANOVA; ns – not significant; *p < 0.05).

**c**. Thin layer chromatography and quantification of cholesterol in rat brain at PND5, PND15 and PND30. n = 8, histograms represent means ± SEM of the measured variable (One-way ANOVA; ns – not significant; **p < 0.01).

**a**

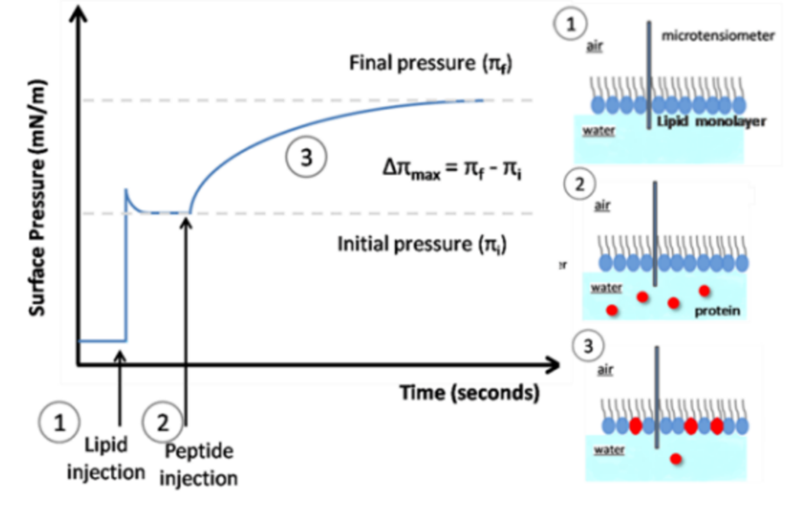


**b**

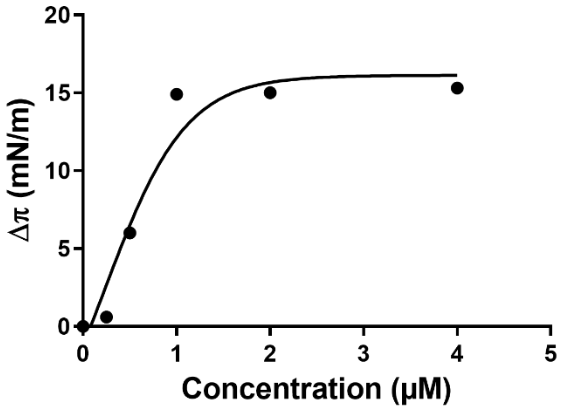


**Figure S5: The interaction between KCC2-GBD and GM1 is saturable.**

**a**. Lipid monolayer principle. 1) Lipids (blue circles) are spread on water drops and the surface pressure of the monolayer is recorded by a microtensiometer. Protein (red circles, in this project WT-KCC2-GBD or W318S-KCC2-GBD) is added into water (2) and its affinity for the lipids allow its interaction and insertion inside the monolayer triggering an increase of surface pressure (3). The difference between initial ($\pi i$) and final ($\pi f$) surface pressure ($\Delta\pi max$) is indicative of the robustness of the interaction.

**b**. Interaction of WT-KCC2-GBD added at different concentrations underneath a GM1 monolayer at an initial pressure of 15 $mN.m^{-1}$. In each case, the surface pressure increase ($\Delta\pi$) was determined after 1 hour of interaction.

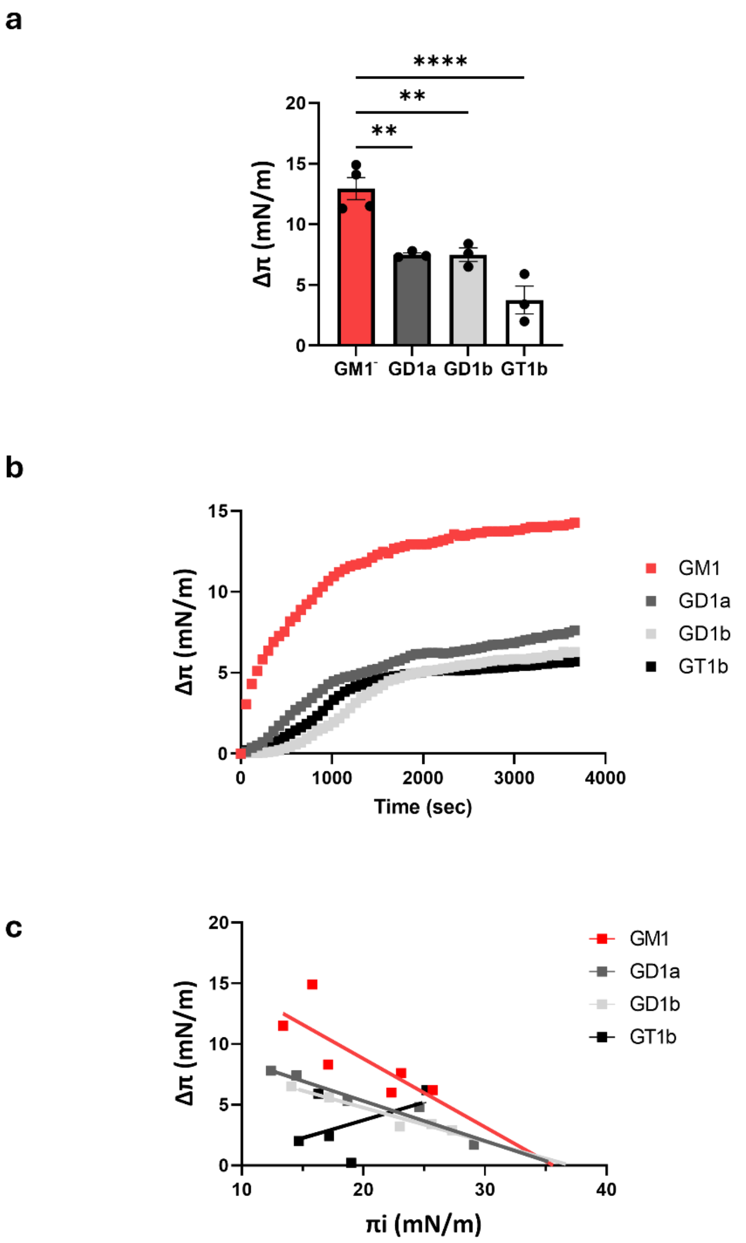


**Figure S6: The KCC2-GBD interacts preferentially with GM1.**
**a**. Structure-activity studies of WT-KCC2-GBD interactions with GM1 (red), GD1a (dark grey), GD1b (red) or GT1b (light grey) (mean ± SEM, n = 3-4, ANOVA test, ****p < 0.0001).
**b.** Kinetics studies of WT-KCC2-GBD interaction with monolayers of GM1 (black), GD1a (dark grey), GD1b (red) or GT1b (light grey). The data show the real-time changes of surface pressure following injection of 10 µM WT-KCC2-GBD beneath the corresponding lipid monolayer prepared at an initial surface pressure of 15 mN·m$^{-1}$. Each curve is representative of three independent experiments.
**c.** Specificity of gangliosides / WT-KCC2-GBD interaction. Neuronal ganglioside monolayers were prepared at various values (initial surface pressure range 10−34 mN·m$^{-1}$). After equilibration of the monolayer, WT-KCC2-GBD (10 µM) was added underneath the GM1 (black), GD1a (dark grey), GD1b (red), or GT1b (light grey) monolayer at a final concentration of 10 µM. The maximal surface pressure increase (Δπmax) was measured after reaching

equilibrium. The critical pressure of insertion was extrapolated as the value of the initial surface pressure at Δπmax = 0.

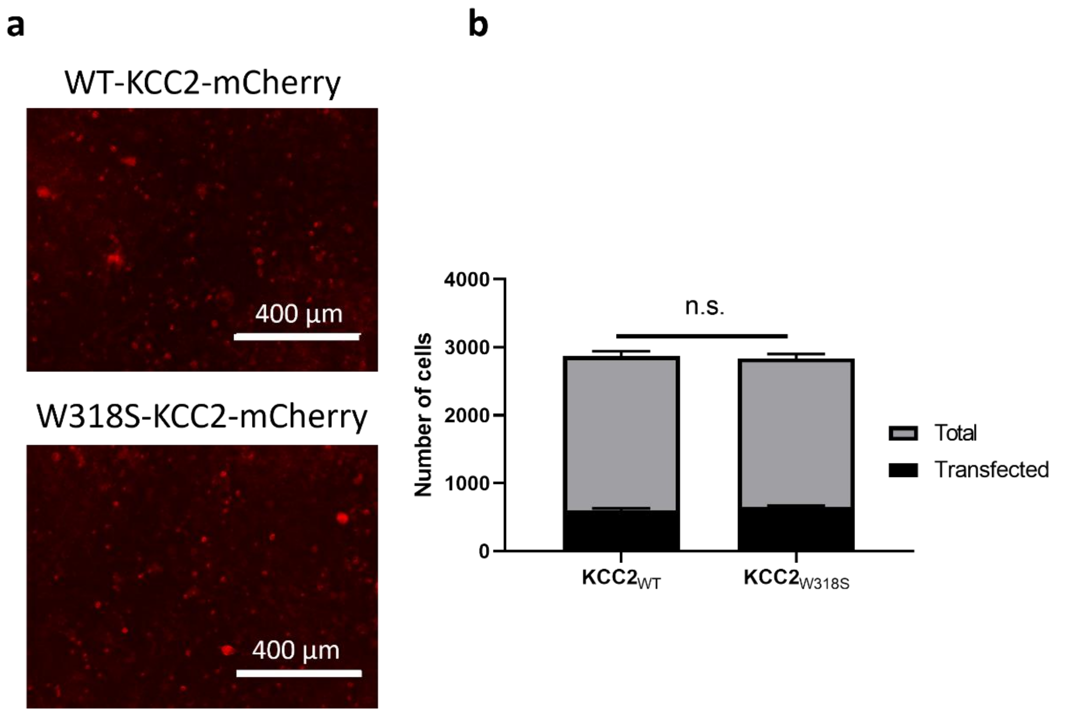


**Figure S7: The W318S-KCC2 mutation does not affect its expression.**

**a**. Representative pictures of transfected HEK293T cells with WT-KCC2-mCherry or W318S-KCC2-mCherry constructs.

**b**. Quantification of the number of transfected cells. Five independent experiments, two-way ANOVA, ns – not significant.

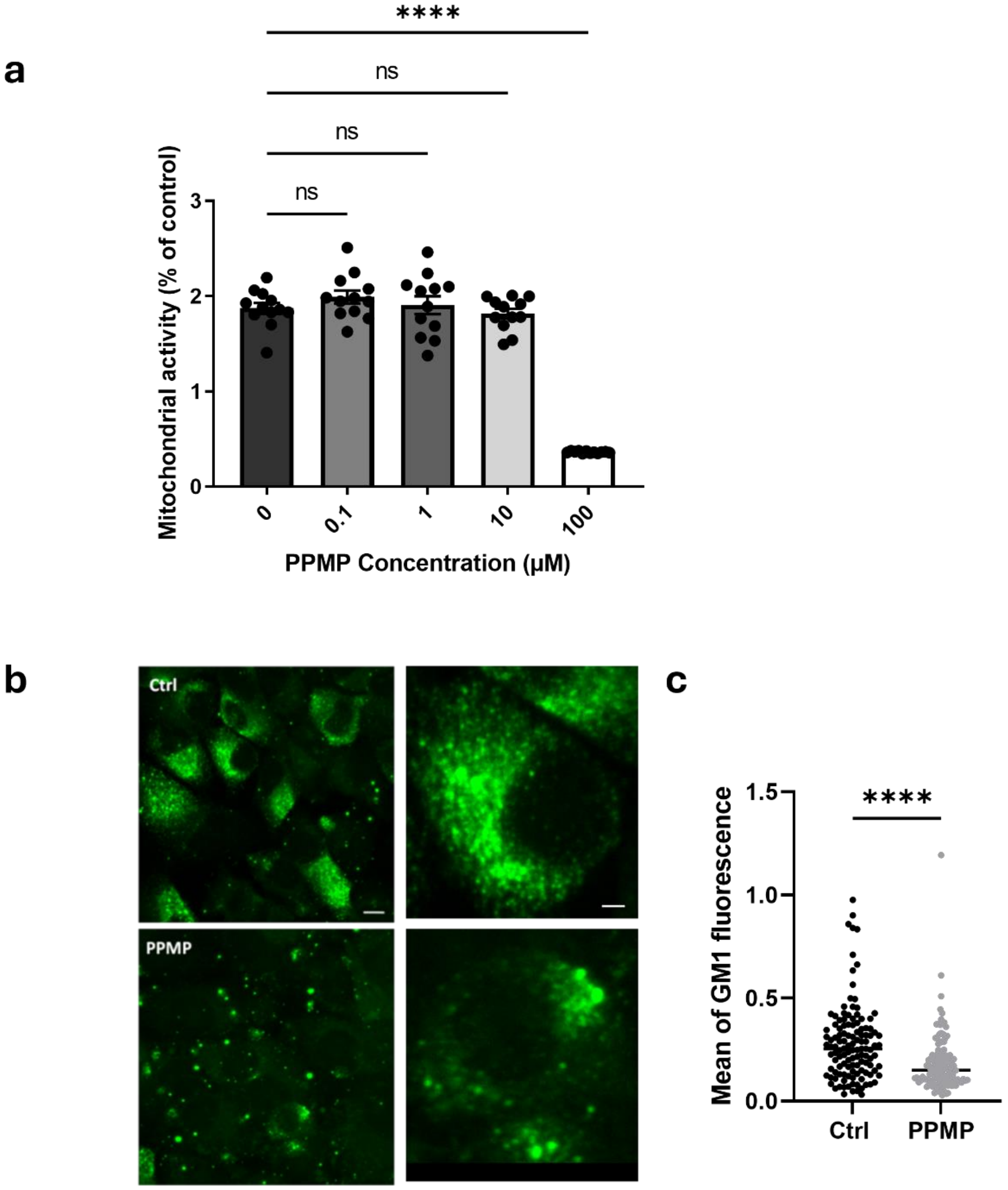


**Figure S8: PPMP treatment decreases GM1 membrane level on HEK293T cells.**
**a.** Mitochondrial activity measurement showing the lack of toxicity of 10 µM PPMP treatment on HEK293 cells. n = 4 independent cultures made in triplicate, mean ± SEM; ANOVA test, ns – not significant; ****p < 0.0001.
**b**. Membrane immunostaining of GM1 on HEK293 cell cultures treated or not with 10 µM PPPM for 48 hours. Pictures are representatives of four independent experiments. Scale bar = 6 µm (left pictures) and scale bar = 2 µm (right pictures).
**c**. Quantification of membrane GM1 immunostaining in control (black) or PPMP-treated HEK293 cells (grey); Mann-Whitney ****p < 0.0001.

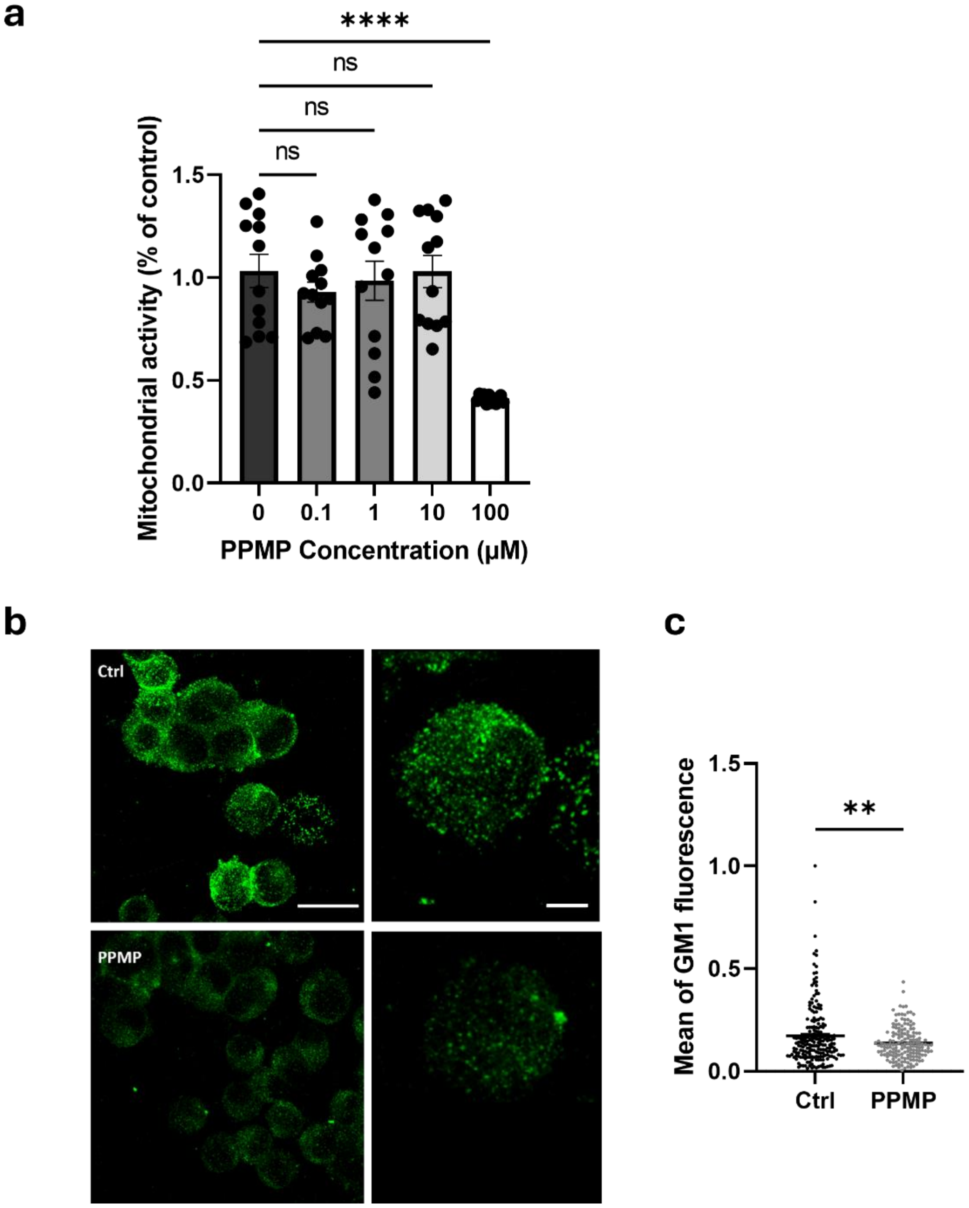


**Figure S9: PPMP treatment decreases GM1 membrane level on Neuro2a cells.**

**a.** Mitochondrial activity measurement showing the lack of toxicity of 10 µM PPMP treatment on Neuro2a cells. N = 4, in triplicate, mean ± SEM; ANOVA test, ns – not significant; ****$p < 0.0001$.

**b**. Membrane immunostaining of GM1 on Neuro2 cell cultures treated or not with 10 µM PPPM for 48 hours. Pictures are representatives of four independent experiments. Scale bar = 25 µm (left pictures) and scale bar = 6 µm (right pictures).

**c**. quantification of GM1 immunostaining in control neurons (black) or PPMP-treated neurons (grey); Mann-Whitney test **$p < 0.01$).

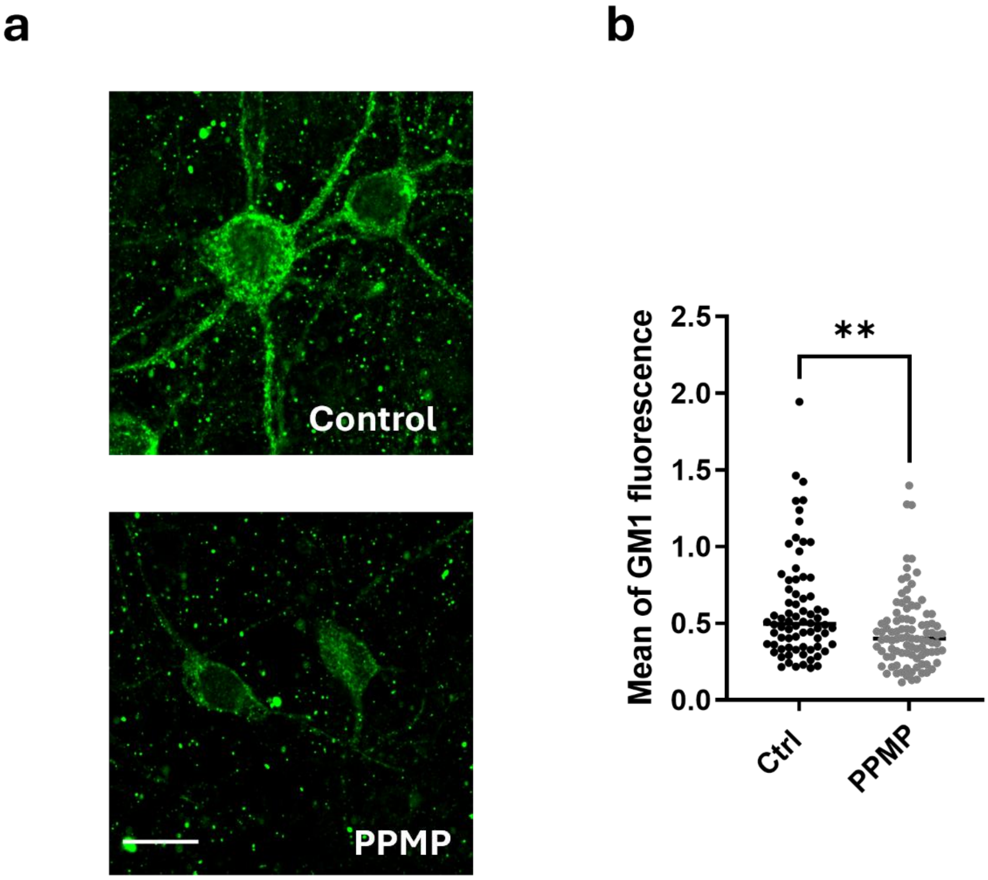


**Figure S10: PPMP treatment decreases GM1 membrane level on hippocampal cultures.**
**a**. Membrane immunostaining of GM1 on primary hippocampal cultures treated or not with 10 µM PPPM for 48 hours. Pictures are representatives of three independent experiments. Scale bar: 15 µm.
**b**. Quantification of GM1 immunostaining in control neurons (black) or PPMP-treated neurons (grey); Mann-Whitney $^{**}p < 0.01$).
